# The ubiquitin conjugase Rad6 mediates ribosome pausing during oxidative stress

**DOI:** 10.1101/2022.09.27.509727

**Authors:** Sezen Meydan, Géssica C. Barros, Vanessa Simões, Lana Harley, Blanche K. Cizubu, Nicholas R. Guydosh, Gustavo M. Silva

## Abstract

Oxidative stress causes K63-linked ubiquitination of ribosomes by the E2 ubiquitin conjugase, Rad6. How Rad6-mediated ubiquitination of ribosomes affects global translation, however, is unclear. We therefore performed Ribo-seq and Disome-seq in *Saccharomyces cerevisiae,* and found that oxidative stress caused ribosome pausing at specific amino acid motifs, and this also led to ribosome collisions. However, these redox pausing signatures were lost in the absence of Rad6 but did not depend on the ribosome-associated quality control (RQC) pathway. We also found that Rad6 is needed to inhibit overall translation in response to oxidative stress and its deletion leads to increased expression of antioxidant genes. Finally, we observed that the lack of Rad6 leads to changes during translation initiation that affect activation of the integrated stress response (ISR) pathway. Our results provide a high-resolution picture of the gene expression changes during oxidative stress and unravel an additional stress response pathway affecting translation elongation.

**HIGHLIGHTS:**

1. Rad6 is required for sequence-specific ribosome pausing under oxidative stress.
2. Rad6 affects translation independently of the RQC pathway.
3. Cells lacking Rad6 show dysregulated translational repression upon oxidative stress.
4. Loss of Rad6 leads to altered activation of the ISR pathway.

## INTRODUCTION

Eukaryotic organisms frequently encounter harmful environmental conditions throughout the course of their lifetime. In these environments, the timely and precise regulation of gene expression is essential to support cellular stress defense, adaptation, and survival (Advani and Ivanov, 2019). Cellular stress caused by the accumulation of reactive oxygen species (ROS) affects important biomolecules such as nucleic acids, proteins, and lipids, and is associated with several pathologies, such as cancer, and cardiovascular and neurodegenerative diseases (Finkel and Holbrook, 2000; Liguori et al., 2018; Sharifi-Rad et al., 2020). ROS, including hydrogen peroxide (H_2_O_2_), can form as a result of metabolic processes but also from exposure to a range of chemicals and pollutants (Picazo and Molin, 2021; Sharifi-Rad et al., 2020). Thus, oxidative stress occurs when ROS production overloads the cellular antioxidant defense. To prevent the detrimental effects of ROS, cells evoke an intricate regulatory network of gene expression both at the transcriptional and translational level (Gasch et al., 2000; Gerashchenko et al., 2012; Picazo and Molin, 2021; Shenton et al., 2006). Although the transcriptional response to oxidative stress has been more extensively studied (Beckhouse et al., 2008; Gasch et al., 2000; Levings et al., 2021; Ma, 2010; Marguerat et al., 2014), a genome-wide understanding of how cells regulate translation in response to oxidative stress is only beginning to be elucidated (Gerashchenko and Gladyshev, 2014; Gerashchenko et al., 2012; Marguerat et al., 2014; Rubio et al., 2021; Shenton et al., 2006).

In response to oxidative stress, eukaryotic cells activate a series of pathways that reprogram translation by shutting down protein production globally while favoring the translation of essential proteins for cell survival (Picazo and Molin, 2021). Some of these pathways are determined by the levels and availability of translation factors or dictated by *cis* regulatory mRNA sequences (Grant, 2011; Picazo and Molin, 2021). However, multiple steps of translation (initiation, elongation, and termination) can be controlled under stress (Grant, 2011), increasing the complexity of the system. Although an extensive number of studies have focused on the regulation of translation initiation (Bresson et al., 2020; Harding et al., 2000; Shenton et al., 2006), mechanisms of elongation regulation during stress are still not well understood.

We previously discovered in the budding yeast *S. cerevisiae* a new mechanism responsible for controlling translation elongation during oxidative stress via ubiquitination of ribosomes (Back et al., 2019; Silva et al., 2015; Zhou et al., 2020). We named this pathway redox control of translation by ubiquitin (RTU) (Dougherty et al., 2020). A key regulator of the RTU is the E2 ubiquitin conjugase Rad6 that rapidly modifies ribosomal proteins with K63-linked polyubiquitin chains in response to H_2_O_2_ (Back et al., 2019; Silva et al., 2015; Simoes et al., 2022; Zhou et al., 2020). Ubiquitinated ribosomes arrest at the pre-translocation stage of translation elongation (Zhou et al., 2020) and are proposed to participate in the shutdown of protein production during stress (Back et al., 2019; Zhou et al., 2020). Furthermore, we recently showed that deletion of *RAD6* prevents K63-linked ubiquitination of ribosomes and leads to continued protein production under oxidative stress and dysregulated levels of antioxidant proteins (Simoes et al., 2022). Rad6 is a multifunctional and highly conserved protein, in which mutations to its human homolog UBE2A are associated with the X-linked intellectual disability type Nascimento (Bruinsma et al., 2016; Nascimento et al., 2006). However, an understanding of the means by which Rad6 controls the translational landscape by modifying ribosomes and the crosstalk of the RTU with other pathways of translation control remains elusive.

To characterize the translational landscape mediated by Rad6 we made use of next generation sequencing of ribosome protected mRNA fragments, also known as Ribo-seq or ribosome profiling. Ribo-seq has uncovered many new aspects of translation dynamics and regulation in a diverse range of organisms and conditions (Ingolia et al., 2009; Ingolia et al., 2019). Moreover, we recently improved our disome profiling (Disome-seq) approach to allow us to identify and quantify the endogenous mRNA sequences occupied by collided ribosomes (disomes) in yeast (Meydan and Guydosh, 2020). Using this method, we were able to show that collisions are widespread events connected with quality control and stress response pathways (Meydan and Guydosh, 2020). We also showed that although most collisions do not activate mRNA decay pathways, they may have an important signaling role in co-translational events (Meydan and Guydosh, 2020). Here, we used both Ribo-seq and Disome-seq combined with RNA-seq to understand the role of Rad6 in translational control during stress. We found that upon hydrogen peroxide treatment, ribosomes from wild-type (WT) cells are preferentially paused on isoleucine-proline enriched sequences. Surprisingly, this redox pausing signature is largely abolished upon deletion of *RAD6*. Furthermore, we showed that the RTU functions independently of the RQC pathway, which is known for detecting and rescuing collided ribosomes. The RQC also relies on K63-linked ubiquitination of ribosomes but deletion of its main E3 ubiquitin ligase *HEL2* did not abolish the burst of K63-linked ubiquitination, or the translation pause signature during stress. Finally, we showed that lack of Rad6 affects translation rates and activates additional translation programs, including the ISR through a non-canonical mechanism. Therefore, this study uncovers a novel mechanism of translational control and positions Rad6 as new remodeler of the translation landscape through a selective ribosome pausing mechanism.

## RESULTS

### Rad6 is required for redox pausing of ribosomes

Rad6-mediated ubiquitination was suggested to affect translation during oxidative stress by arresting translation elongation at the pre-translocation stage (Back et al., 2019; Zhou et al., 2020). To further understand the impact of Rad6-mediated ubiquitination on ribosome pausing at a genome-wide level, we conducted Ribo-seq experiments in wild-type (WT) and *rad6*Δ cells incubated with ± 0.6 mM hydrogen peroxide (H_2_O_2_, “peroxide” hereafter) for 30 min (Figure 1A and S1A) (Ingolia et al., 2009; McGlincy and Ingolia, 2017). This peroxide concentration and the treatment time were optimized based on the peak accumulation of K63-linked polyubiquitin chains after the addition of peroxide to the media (Silva et al., 2015). Supporting the establishment of our system, RNA-seq experiments showed that the peroxide treatment resulted in significantly upregulated expression of genes involved in the oxidative stress response in both WT and *rad6*Δ cells (Figures 1B-C). These results show that both strains respond to this peroxide treatment, allowing us to expand our analysis of the stress response to the translational level.

**Figure 1.**
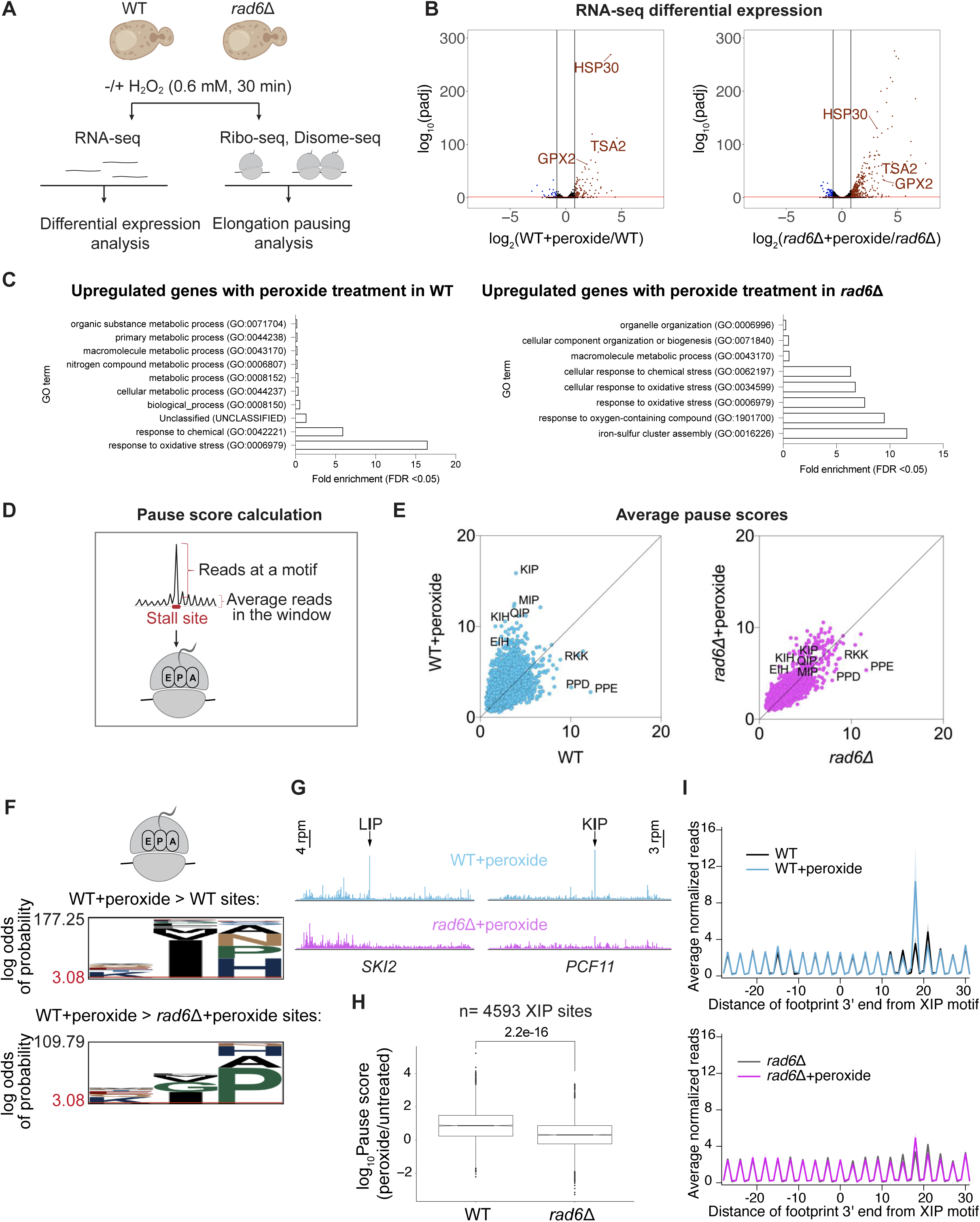
Rad6 is necessary for ribosome pausing during oxidative stress. **(A)** Schematics of RNA-seq, Ribo-seq and Disome-seq experiments conducted in WT and *rad6*Δ *S. cerevisiae* cells ± 0.6 mM H_2_O_2_ (peroxide) for 30 minutes. **(B)** Volcano plot showing the differential RNA expression in WT+peroxide vs WT (left), and *rad6*Δ+peroxide vs *rad6*Δ (right). Genes that are significantly upregulated (log_2_Fold Change > 0.8, padj < 0.05) or downregulated (log_2_Fold Change < -0.8, padj < 0.05), as determined by DESeq2 analysis are shown in red and blue, respectively. The significance cut-off is indicated with a red bar. Known redox gene names are labeled on both graphs. **(C)** Significantly enriched gene ontology (GO) terms of the genes that are upregulated upon peroxide treatment in WT and *rad6*Δ cells. GO analysis is conducted in PANTHER, GO-Slim Biological Process by using *Saccharomyces cerevisiae* (all genes in database) as reference list and test type as FISHER with FDR correction. **(D)** Schematics for calculation of pause score, which is computed by dividing reads at a motif by the average reads in a local window around the motif of interest (± 50 nt). We calculated pause score at tri-amino acid motifs that map within to the ribosomal E (penultimate position of the nascent peptide), P, A sites. **(E)** Average pause scores of 6267 tripeptide motifs plotted for untreated versus peroxide data from WT (left) and *rad6*Δ (right) cells. Each point represents a tripeptide motif. Pause scores were calculated by applying a shift value of 18 nt from the 3’ end of the footprint, placing the first codon of the tripeptide motif in the E site. The average of two replicates is plotted. Some prominent stalling motifs are labeled on the graph. Motifs that correspond to stalling events that appear to increase under peroxide are located above the diagonal. **(F)** pLogo (O’Shea et al., 2013) motif analysis of the tri-amino acid motifs that have 1.5 fold higher average pause score in WT+peroxide vs WT samples (top, n_foreground_=1004, n_background_=6996) and in WT+peroxide vs *rad6*Δ+peroxide (bottom, n_foreground_=397, n_background_=7603) samples. The plots show enrichment for motifs with Ile in P site and Pro in the A site for WT+peroxide samples vs WT or *rad6*Δ+peroxide cells. **(G)** Example individual genes (*SKI2* and *PCF11),* where the effect of Rad6 on redox pausing is observed. The data are obtained from pooled biological duplicates. The stalling peaks corresponding to LIP and KIP motifs are indicated by arrows (top, blue traces) and are lost in the absence of Rad6 (bottom, pink traces). **(H)** Box plot showing the significant loss of redox pausing at XIP sites (n=4593) in *rad6*Δ cells compared to WT. The significance of differences in the median of these individual pause scores were computed by independent 2-group Mann Whitney U Test. **(I)** Average normalized Ribo-seq rpm mapped to genes aligned by their respective XIP motifs in WT (top) and *rad6*Δ (bottom) cells show loss of redox pausing at these motifs on average (top blue trace vs bottom pink trace). Standard deviation of two replicates is shown by shaded error bar.

To understand the effect of Rad6 on translation elongation and to globally quantify ribosome pausing differences, we computed “average pause scores” for every combination of tri-amino acid motif (Figure 1D and Methods). First, we observed that amino-acid sequences such as PPD, PPE and RKK caused the strongest pausing in untreated WT cells (Figure 1E, left panel). However, the relative level of pausing at these sequences was not as high as in stressed cells. Instead, we found that peroxide treatment in WT cells caused reprogramming of ribosome arrest and resulted in elevated pausing at specific sequence motifs, which we define as “redox pausing” (Figure 1E, left panel). These redox pausing signatures were enriched in sites that have proline (Pro) and histidine (His) codons at the ribosomal A-site. We also observed that stalling at isoleucine (Ile) codons at the ribosomal P-sites increased upon peroxide treatment, especially in combination with A-site Pro codons (Figures 1E-F). Since we did not detect substantial enrichment for residues that specifically mapped to the E-site (*i.e.* the amino acid corresponding to the penultimate, C-terminal position of the nascent peptide), we designated this stalling motif as “XIP”, where X refers to any amino acid composition.

Given prior evidence that Rad6-mediated ubiquitination could modulate translation elongation (Simoes et al., 2022), we hypothesized that loss of accumulated ubiquitination by Rad6 under oxidative stress could affect ribosome stalling. To test this hypothesis, we performed our pausing analysis in *rad6*Δ cells. Strikingly, we found that redox pausing signatures were lost in cells lacking Rad6 (Figure 1E, right panel). The previously identified XIP redox pausing motifs were the most susceptible to the loss of Rad6 (Figure 1F). Consistently, analysis of individual or averaged ribosome occupancy at XIP sites also revealed considerable loss of redox pausing in the absence of Rad6 (Figures 1G-H). In the untreated cells, ribosome stalling signatures looked similar between the two strains except for increased stalling in *rad6*Δ cells at A-site Trp codons, a result that is specific to the SUB280 background used here (Figure S1B, see below for further discussion). As expected, plots of average ribosome occupancy centered at XIP motifs were consistent with the pause score analysis and revealed a peroxide-induced stalling peak that was absent in untreated WT cells and in the *rad6τι* strain (Figure 1I, top and bottom panel, respectively).

Upon arresting at XIP motifs, ribosomes could resume translation or be rescued by quality control systems to make them available for new rounds of translation. To determine whether these XIP motifs lead to ribosome rescue, we developed an inducible dual luciferase reporter, in which we can insert sequences between Renilla Luciferase (Rluc) and Firefly Luciferase (Fluc) coding regions that are expected to stall translation and lead to ribosome rescue. As a proof of principle, we determined that several sequences identified with a high pause score in our Ribo-seq data indeed prevent the synthesis of Fluc (Figure S1C). Because of the global translation repression that occurs under oxidative stress (Figure S1D), this method did not allow us to measure dynamic changes in the Fluc/Rluc ratios during the short time window following peroxide treatment used in this study. However, we still observed a reduction in the Fluc/Rluc ratio when 3xKIP was inserted in the absence of peroxide, consistent with an interpretation that there could be a loss of ribosomes between Fluc and Rluc in WT cells due to rescue or drop off. This trend was unchanged in *rad6*Δ cells, suggesting that Rad6 does not promote ribosome rescue (Figure 2A).

**Figure 2.**
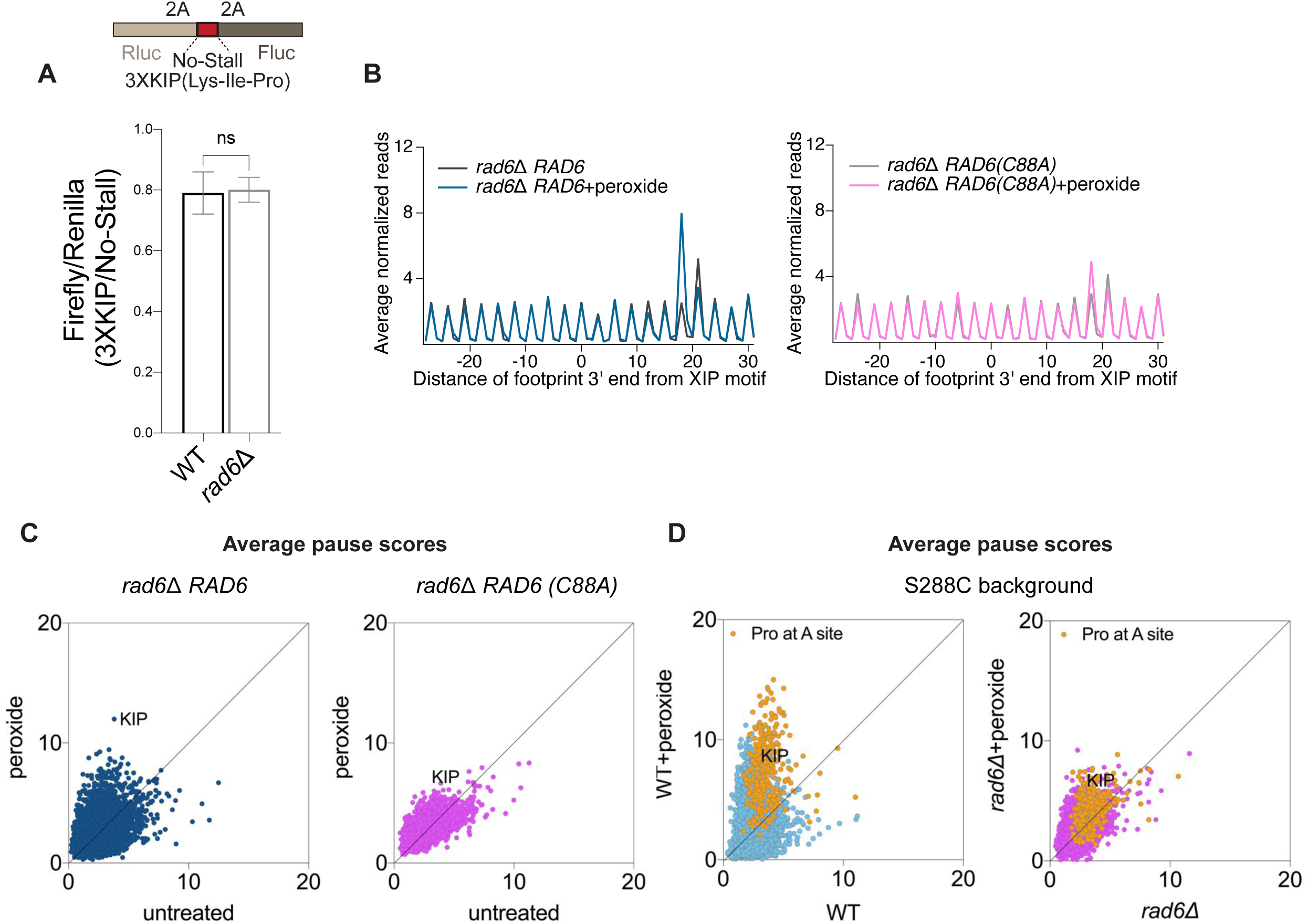
Ubiquitination activity of Rad6 mediates ribosome pausing but not rescue. **(A)** Schematics for the Renilla-Firefly reporter construct used for the experiments to measure ribosome rescue at a redox pausing motif (top, 3XKIP). The Fluc/Rluc ratio is expected to become lower when ribosomes dissociate from the mRNA (i.e. via ribosome rescue) after translating the Rluc sequence but prior to reaching the Fluc sequence. The ratio of the Fluc/Rluc value for the 3XKIP reporter compared to a No-Stall reporter is shown (bottom). Note that the observed value of ∼0.8 indicates some level of ribosome rescue due to the 3XKIP motif. Deletion of *RAD6* does not appear to affect ribosome rescue in this case. The significance is assessed by one-way Anova test. ns = not significant. **(B)** Average normalized Ribo-seq rpm mapped to genes aligned by their respective XIP motifs in *rad6*Δ cells complemented with either WT Rad6 (*rad6*Δ *RAD6,* left) or its catalytically dead mutant (*rad6*Δ *RAD6(C88A)*, right) show that ubiquitination activity of Rad6 is necessary to restore redox pausing at XIP. **(C)** Average pause scores of 6267 tripeptide motifs plotted for untreated versus peroxide data from *rad6*Δ *RAD6* (left) and *rad6*Δ *RAD6 (C88A)* cells (right) show that expression of WT Rad6 restores overall redox pausing but its catalytically dead mutant cannot. The KIP motif is labeled. **(D)** Average pause scores of 6267 tripeptide motifs plotted for untreated versus peroxide data from S288C WT and *rad6*Δ cells. Motifs with Pro codons at A site indicated in yellow and KIP motif labeled. These data indicate that the redox pausing signatures and effect of Rad6 loss on them are consistent between different yeast strains.

Because oxidative stress induced by peroxide was previously associated with increased translation of 5’ and 3’ untranslated regions (UTRs) (Gerashchenko and Gladyshev, 2014; Gerashchenko et al., 2012), we also tested whether Rad6 impacts these translation events outside of main open reading frames. Metagene analysis, performed by averaging data from genes aligned by their start or stop codons, showed modest changes in the occupancy of 5’ and 3’ UTRs with peroxide treatment in WT cells, as expected (Figures S1E-F). The absence of Rad6 did not affect these trends (Figures S1E-F), suggesting that Rad6 does not strongly impact translation of UTR regions under oxidative stress but mainly affects elongation in the main ORF.

To confirm our observation that Rad6 has a specific role in modulating redox pausing, we conducted additional Ribo-seq experiments in which we expressed both the WT Rad6 and the catalytically dead mutant Rad6 (Rad6^C88A^) in a *rad6*Δ background. Expression of WT Rad6 in *rad6*Δ cells restored peroxide-induced stalling at the XIP motif and other sites, whereas the ubiquitination-deficient mutant Rad6^C88A^ did not (Figures 2B-C). These results suggest that Rad6 catalytic activity is essential to regulate redox pausing.

To confirm the generality of our results, we also considered the effect of the yeast strain background. The yeast strains used in this study (SUB280 background) were constructed to express a single ubiquitin gene episomally (Finley et al., 1994). To test whether the unique properties of this strain were related to the observed redox pausing signatures, we repeated experiments in the S288C background. Experiments in S288C did not show an increase in A-site pausing at Trp codons in the *rad6*Δ cells (Figure S1B), suggesting that this particular effect is not a robust feature of redox pausing. However, they did recapitulate the previously observed redox pausing signatures at A-site Pro codons, including XIP motifs (Figure 2D). These data therefore show that redox pausing is a consistent mechanism of translational control in response to stress and that Rad6 plays a key role in this translation phenotype.

### Redox pausing signatures are not mediated by the RQC pathway

We next investigated whether Rad6 activity plays any role in the well-established RQC pathway. Stalled ribosomes can physically block upstream ribosomes from translating, resulting in the formation of a ribosome collision complex called a disome, where the two ribosomes interact. The RQC pathway is a cellular mechanism that detects disomes and promotes their removal from mRNAs. Therefore, we first evaluated whether peroxide treatment would produce disomes that could be rescued by the RQC. We, and others, have previously established the Disome-seq technique in yeast to find the genome-wide distribution of collided ribosomes in the cell (Guydosh and Green, 2014; Meydan and Guydosh, 2020; Zhao et al., 2021). Disome-seq in WT cells also showed evidence of redox pausing signatures, such as the XIP motif and generally mirrored our Ribo-seq data (Figure S2A-B). This suggests that stalled ribosomes formed during oxidative stress also collide with each other.

Previous studies showed that an E3 ubiquitin ligase, Hel2, triggers the RQC pathway by ubiquitinating collided ribosomes stalled at positively charged amino acid sequences, such as poly-Arg or poly-Lys (Brandman et al., 2012; Houston et al., 2022; Ikeuchi et al., 2019; Matsuo et al., 2017; Matsuo et al., 2020). Ubiquitination leads to ribosome rescue but, in the absence of Hel2, collided ribosomes bypass these stall-inducing sequences and continue translating. Since the RQC pathway regulates translation arrest via Hel2-mediated ubiquitination of ribosomes, it raises the question of whether the E3 Hel2 and E2 Rad6 cooperate in the same pathway of translational control.

To further investigate whether Hel2 would be involved in rescuing disomes formed in response to stress, we performed a Ribo-seq experiment in *hel2*11 cells. During peroxide treatment, XIP redox pausing signatures were still present in cells lacking Hel2, which suggests a separation of functions (Figure 3A). Consistent with the idea that Hel2 does not work with Rad6 to regulate translation under oxidative stress, we also showed that loss of Hel2 did not affect the burst of K63 ubiquitination induced by peroxide treatment (Figure 3B), further supporting that Rad6 activity is independent of Hel2 and the RQC. To explore the activity of Hel2 and Rad6 in rescuing stalled ribosomes, we inserted a known RQC-targeted stalling sequence consisting of 6 consecutive Arg codons (6XCGA) into our Renilla-Firefly luciferase reporter for ribosome rescue. This sequence is particularly problematic for the ribosome to translate due to I-C wobble codon-anticodon pairing (Letzring et al., 2013). Ribosomes stalled at 6XCGA are known to be rescued by RQC (Letzring et al., 2013), and as expected, we observed more ribosomes bypassing this stall-inducing site in the absence of Hel2 (Figures 3C and S2C). However, deletion of *RAD6* did not result in a significant increase in Fluc/Rluc signal of the 6XCGA reporter (Figure 3C and S2C). These results suggest that Rad6 does not influence ribosome stalling and rescue in the same way as Hel2. Collectively, our findings further indicate that Hel2 operates on a subpopulation of arrested ribosomes that likely does not include those (i.e. XIP motifs) that are enhanced by oxidative stress.

**Figure 3.**
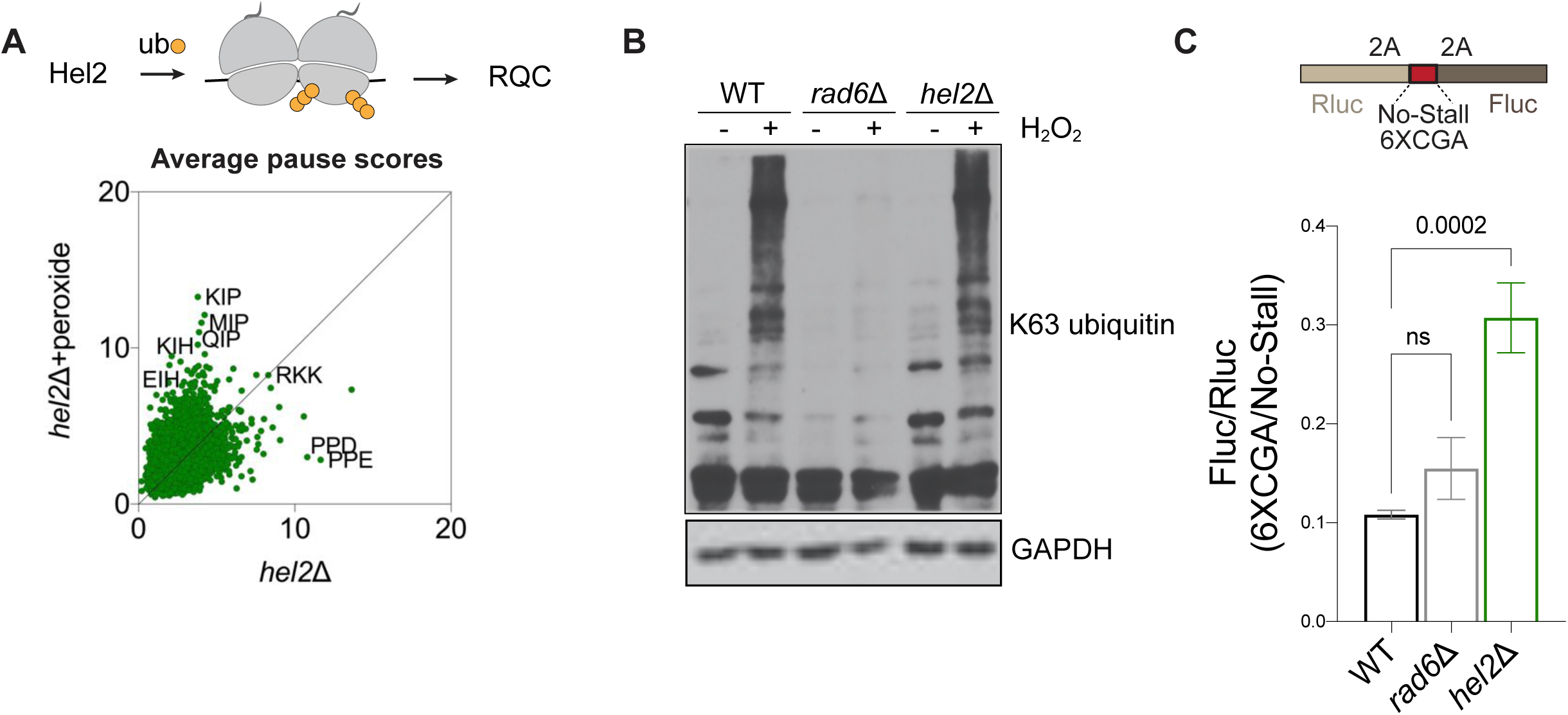
Redox pausing is not mediated by the RQC pathway. **(A)** Hel2 is a K63 ubiquitin ligase that detects disomes and triggers the RQC pathway (top). Average pause scores of 6267 tripeptide motifs plotted for untreated versus peroxide data from *hel2*11 cells show that redox pausing signatures are intact in the absence of Hel2 (bottom). **(B)** Western blotting demonstrates that deletion of *RAD6* eliminates peroxide induced K63 ubiquitination whereas *hel2*11 does not affect it. GAPDH is used as a loading control. **(C)** The schematics of the Renilla-Firefly construct used for the reporter experiments to measure ribosome rescue of an RQC-targeting sequence (top, 6XCGA). The Fluc/Rluc ratio of the 6XCGA reporter compared to No-Stall reporter is shown (bottom). Deletion of *HEL2* causes increased Fluc/Rluc since ribosomes are no longer rescued, whereas deletion of *RAD6* does not significantly affect ribosome rescue at this sequence. The significance is assessed by one-way Anova test. ns = not significant.

### Rad6 is required for translational repression during oxidative stress

Having established that deletion of Rad6 affects ribosome stalling in a way that is different from Hel2, we further explored mechanisms that could be responsible for the redox-induced pausing of ribosomes. We first hypothesized that differential availability of tRNAs could stall ribosomes at XIP motifs during peroxide treatment. It was previously shown that peroxide treatment causes degradation of Pro-tRNA^AGG^, resulting in a ribosome stalled with an empty A-site as it waits for binding of prolyl-tRNA (Wu et al., 2019). One possibility for the loss of redox pausing in *rad6*Δ cells could be that Rad6 indirectly or directly mediates the degradation of prolyl-tRNAs. Loss of Rad6 then might stabilize prolyl-tRNAs and thereby alleviate ribosome stalling at Pro codons. To test this possibility, we conducted northern blotting experiments with a probe that matched Pro-tRNA^AGG^. We found that the peroxide concentration that we used for our Ribo-seq, Disome-seq and RNA-seq experiments (0.6 mM) did not result in tRNA degradation (Figure 4A, lane 2). Only at higher concentrations of peroxide (9.8 mM, as used in (Wu et al., 2019), and 98 mM), we were able to detect the appearance of a tRNA fragment (Figure 4A, lanes 3-4, indicated by an arrow). We also observed that the magnitude and concentration dependence of prolyl-tRNA degradation induced by peroxide was the same in the *rad6*Δ strain (Figure 4A, lanes 5-8), further suggesting that loss of redox pausing in the absence of Rad6 is not due to changes in tRNA stability.

**Figure 4.**
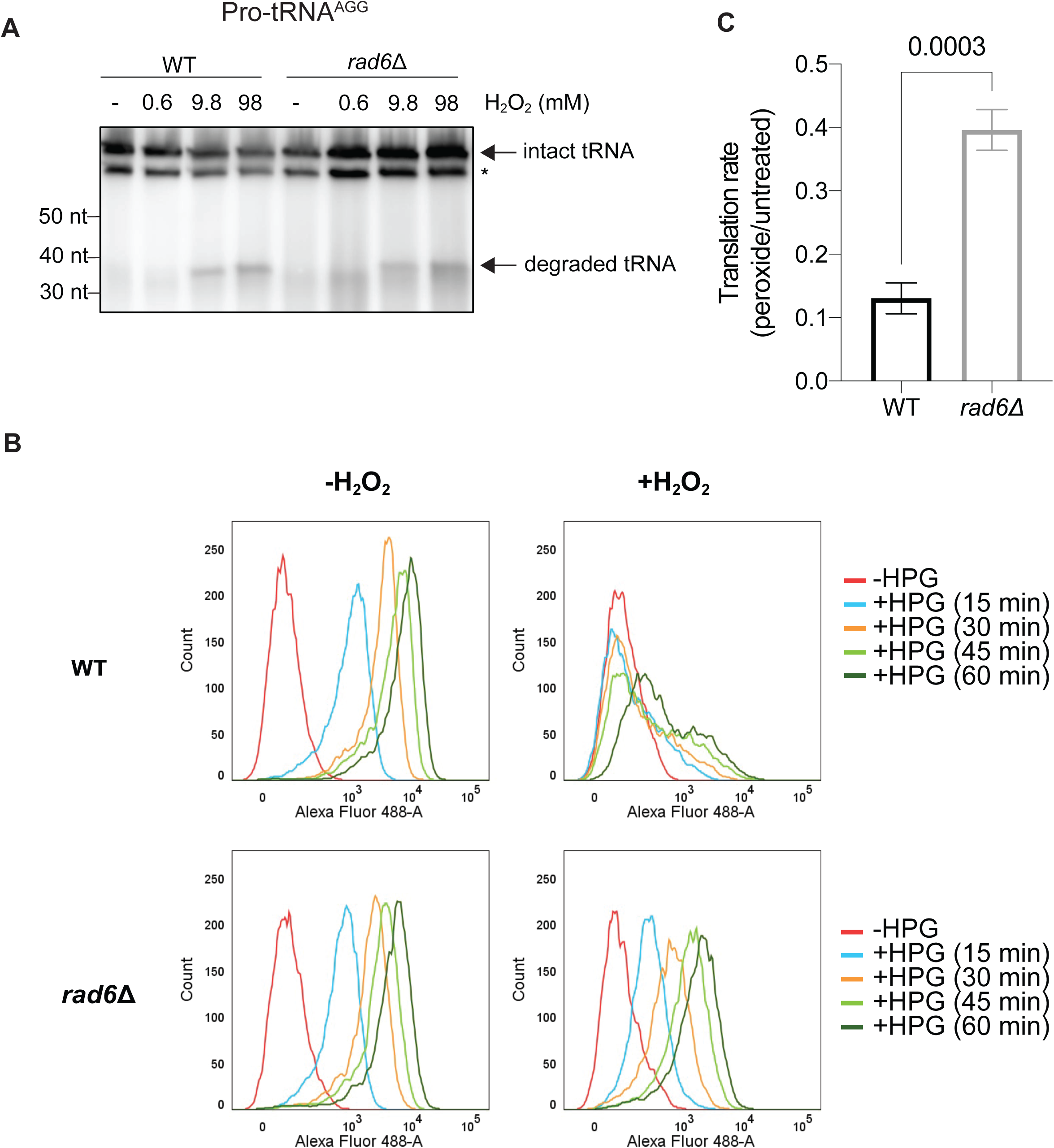
Rad6 promotes translation inhibition during oxidative stress. **(A)** Northern blotting by using a probe to Pro-tRNA^AGG^ shows that deletion of Rad6 does not affect peroxide induced degradation of this tRNA, which only takes place at very high concentrations of peroxide. The degradation fragment and intact tRNA are indicated by arrows. Asterisk (*) refers to a non-specific band. **(B)** Histograms of the HPG incorporation assay showing the number of cells (y-axis) and fluorescence magnitude (x-axis) at indicated time points of HPG incubation (15-60 minutes). WT cells exhibit decreased HPG incorporation in the presence of peroxide (top panel) and this inhibition is slowly released over time. In contrast, HPG incorporation in *rad6*Δ cells is affected less by the peroxide treatment (bottom panel). **(C)** Quantification of HPG incorporation during peroxide treatment is shown as a normalized rate for HPG incorporation in treated vs untreated cells. The translation rates were calculated by fitting the mean fluorescence values to a linear regression as a function of time. Significance is determined by two-tailed unpaired t-test.

We next assessed the overall rate of translation in *rad6*Δ cells to see if the absence of redox pausing was related to the cell’s ability to make proteins. We evaluated changes in translation rates by incorporation of a methionine analog, Homopropargylglycine (HPG). This assay captures the totality of all effects on translation, including changes to both initiation and elongation, and provides an opportunity to evaluate mechanistic models. One possibility is that in the absence of Rad6, ribosomes would no longer undergo redox pausing and could therefore generate more protein during stress. Consistent with this model, the drop in the rate of translation in *rad6*Δ cells due to oxidative stress was significantly less than in WT cells (Figures 4B-C, S3A). While translation in WT cells was severely inhibited by peroxide treatment, cells lacking Rad6 were significantly less sensitive to this translation repression. In addition, the effect of Rad6 on peroxide-induced translational repression was reproducible in the S288C background (Figure S3B-C). These findings are also in agreement with previous data showing higher puromycin incorporation in *rad6*Δ cells in the presence of peroxide compared to WT cells (Simoes et al., 2022). Overall, our results show that Rad6 is necessary for the oxidative stress-induced translational repression.

### Rad6 activity induces eIF2α phosphorylation

To understand the physiological impact of dysregulated translation in the absence of Rad6, we next explored by RNA-seq how peroxide affects the transcriptome in WT and *rad6*Δ cells. In the absence of oxidative stress (untreated), *rad6*Δ cells had significantly upregulated expression of multiple genes involved in metabolic processes (such as *GDB1, GPH1, PKP1, DAK2*), the heat shock response (such as *HSP26*, *HSP30*, *HSP78*), and oxidative stress response (such as *GRX1, SOD2, TSA2,* PRX1) compared to WT cells (Figure S4A). This suggests that even without oxidative stress, the lack of Rad6 causes a mild stress response. We next looked at the gene expression changes upon peroxide treatment. In *rad6*Δ cells, we found that the expression of oxidative stress response genes is upregulated beyond that found in the equivalently treated WT cells (Figures 1B, 5A and S4B), consistent with the previous observation of increased ROS in *rad6*Δ cells under peroxide (Simoes et al., 2022). When we limited our analysis to 21 genes coding for known antioxidant enzymes, we observed overactivated expression of these redox genes in *rad6*Δ cells in both untreated and peroxide-treated samples (Figures 5B and S4B). In addition, there were fewer ribosomal protein transcripts in *rad6*Δ cells, and this downregulation was exacerbated with peroxide treatment (Figure S4C). Decreased level of ribosomal protein transcripts is a hallmark of TOR inactivation, which is likely being driven by ROS in these cells (Picazo and Molin, 2021). We also observed that the translation efficiency of redox genes (ribosome footprints per transcript) was significantly increased by loss of Rad6 in peroxide-treated cells (Figure S4D). These results support a model that in the absence of Rad6, the increased ROS in the cell drives a distinct response at the RNA and translational level.

**Figure 5.**
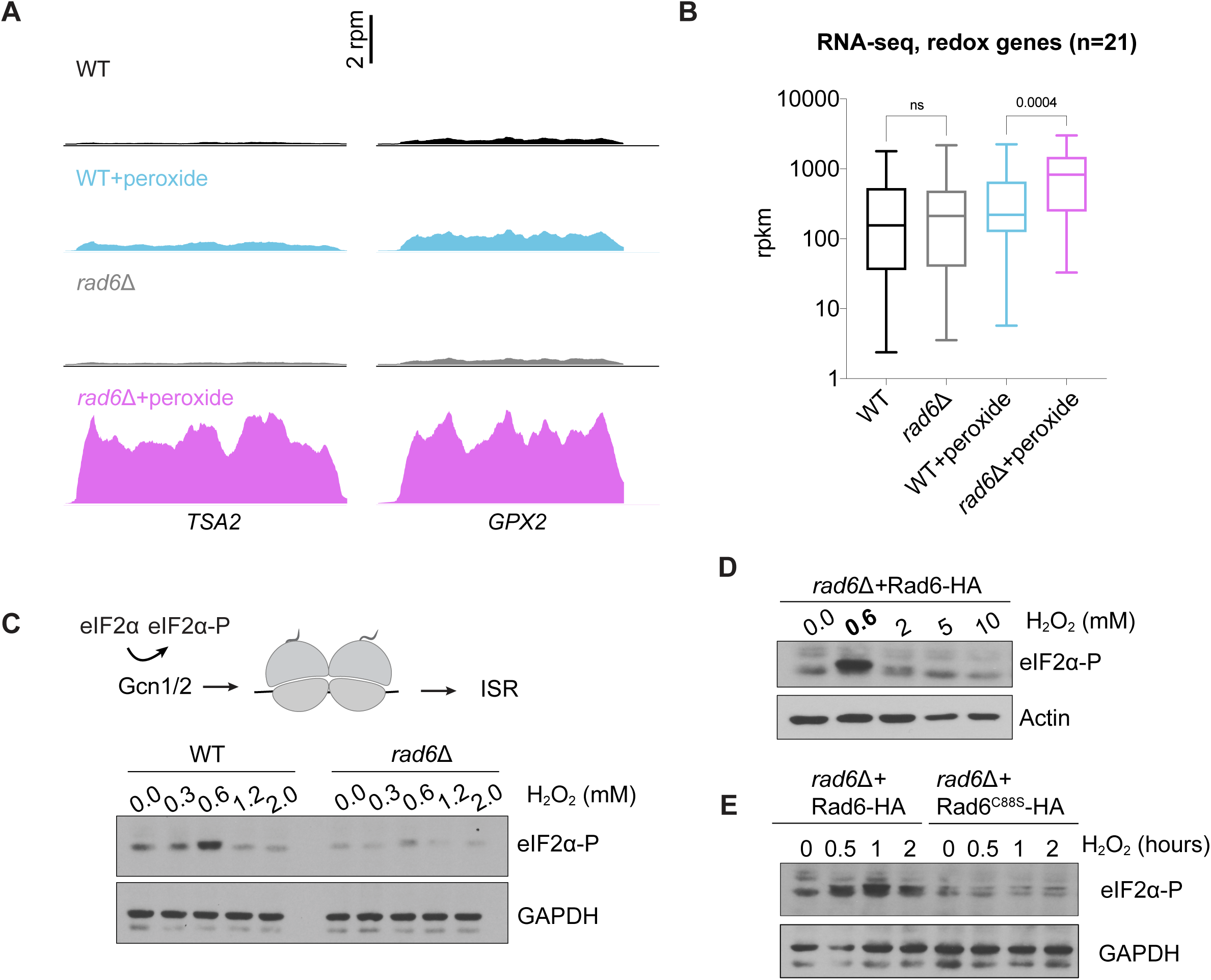
Rad6 promotes eIF2α phosphorylation. **(A)** RNA-seq snapshots of *TSA2* and *GPX2* genes show that the lack of Rad6 causes increased expression of these mRNAs during peroxide treatment (pink versus blue traces). The data is obtained from pooled biological duplicates. **(B)** RNA-seq levels for the mRNAs encoding 21 redox enzymes show upregulation of these genes in *rad6*Δ cells under peroxide treatment. Significance is calculated by one-way Anova test. ns = not significant. **(C)** Disome detection by Gcn1/2 induces eIF2α phosphorylation and ISR activation. Western blotting demonstrates that eIF2α is phosphorylated upon oxidative stress, reaching a maximum at 0.6 mM peroxide, suggestive of increased ribosome stalling events. Peroxide-induced eIF2α-P is reduced in the absence of Rad6, consistent with less ribosome stalling. GAPDH is used as a loading control. **(D)** Western blotting shows that the *rad6*Δ cells complemented with WT Rad6 (Rad6-HA) restores peroxide-induced eIF2α phosphorylation. Actin is used as a loading control. **(E)** Western blotting shows that the *rad6*Δ cells complemented with WT Rad6 (Rad6-HA) restores peroxide-induced eIF2α phosphorylation but the catalytically dead mutant (Rad6^C88S^-HA) cannot, even at longer incubation times with 0. 6 mM peroxide. GAPDH is used as a loading control.

Since *rad6*Δ cells seem to display a higher basal level of stress, we reasoned that loss of Rad6 could lead to the specific activation of the ISR pathway (also known as general amino acid control pathway in yeast) that is known to be induced by many stresses, including peroxide treatment (Grant, 2011; Picazo and Molin, 2021). Oxidative stress-induced ISR results in phosphorylation of eIF2α (eIF2α-P) by the Gcn2 kinase and its coactivators Gcn1 and Gcn20 (Mascarenhas et al., 2008). This leads to repression of overall translation while activating the transcription of stress response genes. In WT cells, we observed the expected induction of the ISR (increased eIF2α-P) during peroxide treatment, reaching its maximum level at 0.6 mM (Figure 5C). Surprisingly, however, phosphorylation of eIF2α in *rad6*Δ cells remained low in response to oxidative stress (Figures 5C and S4E, and Discussion). Expression of Rad6^WT^, but not its ubiquitination deficient mutant (Rad6^C88S^), in *rad6*Δ cells restored eIF2α-P at 0.6 mM peroxide (Figure 5D). Even longer incubation times with peroxide did not result in increased levels of eIF2α-P in *rad6*Δ cells expressing the mutant Rad6^C88S^ (Figure 5E). These observations could be explained by a model where lack of Rad6 affects the expression of the ISR machinery and thereby impairs regulation of eIF2α phosphorylation. To test this possibility, we checked if the components of the ISR machinery, Gcn1, Gcn2 and Gcn20, are properly expressed in the absence of Rad6. The Ribo-seq and RNA-seq data showed no difference in RNA levels and translation efficiency (TE) of the genes coding for Gcn1/2/20 proteins (Figure S4F), suggesting that the absence of Rad6 does not directly affect the expression of ISR machinery components. Another possibility is that loss of Rad6 activates the TOR pathway, which in turn inhibits Gcn2 activity (Cherkasova and Hinnebusch, 2003; Kubota et al., 2003). However, our RNA-seq data showed that the TOR pathway is likely inhibited in *rad6*Δ cells (Figure S4C), suggesting that loss of ISR activation in *rad6*Δ cells is not due to TOR activation.

### Lack of Rad6 confers constitutive *GCN4* translation

Since we observed a lack of peroxide-induced eIF2α-P in *rad6*Δ cells (Figure 5C), we expected the ISR, and its associated effects on the translation of the *GCN4* gene, would not occur. However, we noticed that translation of the *GCN4* gene was increased in this strain (while transcript level remained constant) and this was true both in the presence and absence of peroxide (Figure 6A). This is an unexpected effect because the accumulation of eIF2α-P is the typical driver of increased translation of the *GCN4* mRNA, which encodes a transcription factor that upregulates many ISR genes (Hinnebusch, 2005).

**Figure 6.**
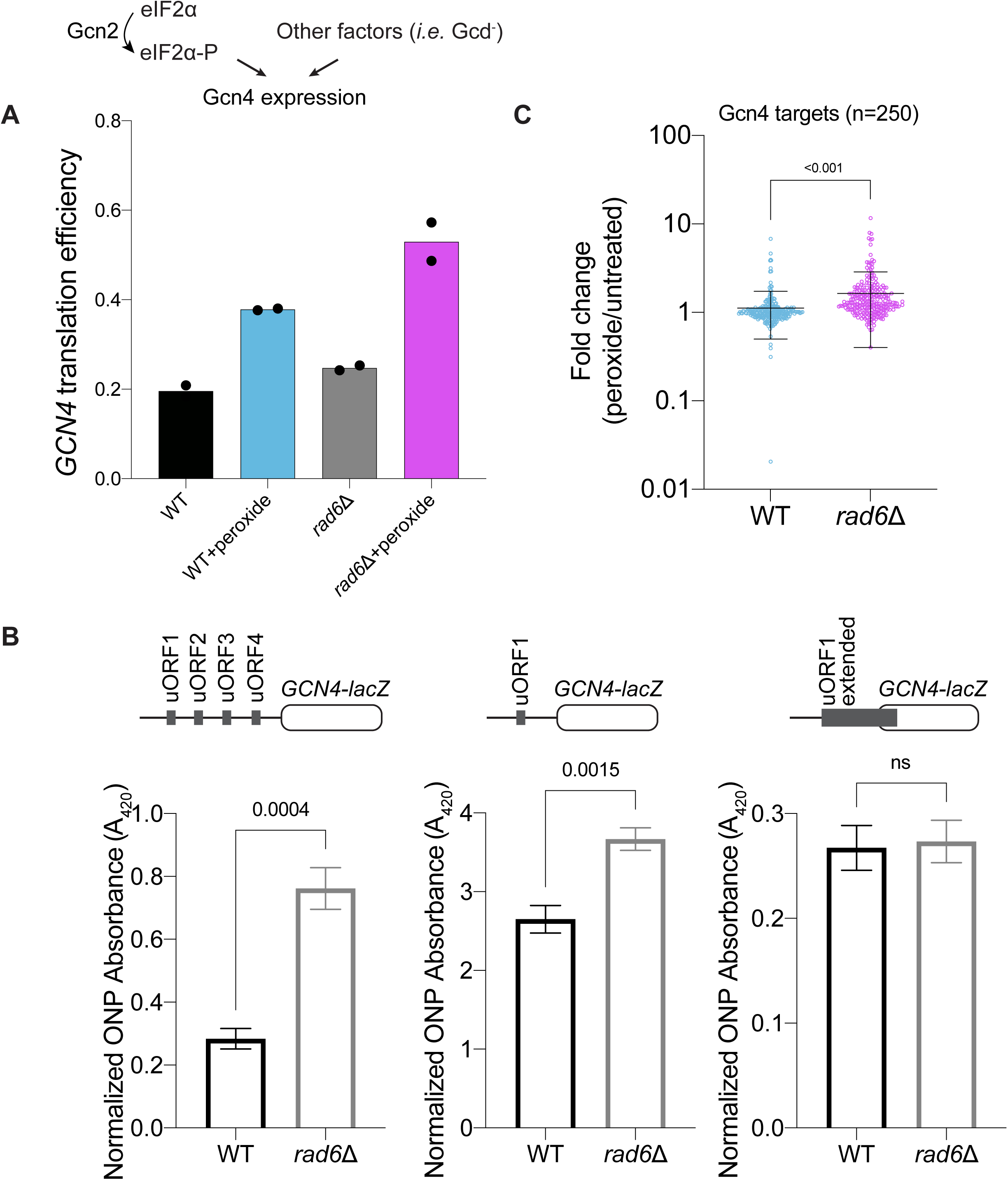
Lack of Rad6 leads to constitutive translation of Gcn4. **(A)** eIF2α phosphorylation and other events (such as loss of initiation factors referred to as “Gcd^-^” phenotype) induce translation of Gcn4. Bar chart shows that *GCN4* translation efficiency (Ribo-seq reads normalized to RNA-seq reads for *GCN4* main ORF) increases with peroxide in both WT and *rad6*Δ cells. Translation efficiency of *GCN4* is higher in *rad6*Δ cells. **(B)** Reporter assay for *GCN4* activation. *GCN4* activation is assayed by *lacZ*, which is fused to the coding sequence. At top, constructs used are shown: all 4 uORFs (left, main construct to measure Gcn4 translation with all native uORFs present), with only uORF1 (middle, as a positive control for activation) and an extended version of uORF1 (right, to assess leaky scanning). The y-axes show ONPG absorption values at 420 nm, normalized by total protein levels. The statistical significance is calculated by unpaired t-test. The data show that loss of Rad6 increases *GCN4* translation and this effect is not due to leaky scanning. **(C)** Peroxide-induced expression of Gcn4’s transcriptional targets (n=250, see Methods for further details) assessed by RNA-seq shows that constitutive translation of Gcn4 in *rad6*Δ cells also leads to increased expression of its downstream genes.

We therefore explored alternative mechanisms that could increase translation of *GCN4*. The *GCN4* gene has 4 upstream open reading frames (uORFs) (Figure 6B). After translating uORF1, many ribosomes are not fully recycled and the 40S subunits remain on the mRNA (Hinnebusch, 2005). These 40S subunits then resume scanning and rebind the ternary complex (eIF2-GTP-tRNAi-Met), which allows them to reinitiate translation at uORF2, uORF3, or uORF4. Termination and recycling after these uORFs are generally more efficient, which prevents the ribosomes from reinitiating again and translating the *GCN4* main ORF. During stress conditions, eIF2α-P reduces ternary complex (TC) levels, and thereby increases the odds that 40S subunits bypass uORF2, uORF3 and uORF4 and instead reinitiate translation at the *GCN4* main ORF. In addition, leaky scanning, where ribosomes skip the uORFs and instead initiate at the downstream *GCN4* main ORF, or other initiation defects can also lead to increased *GCN4* translation. Since eIF2α is not phosphorylated in *rad6*Δ cells with oxidative stress, the observed high translation efficiency of *GCN4* could be due either due to initiation defects or leaky scanning. To monitor *GCN4* main ORF expression in relationship to its uORFs, we used a *GCN4-lacZ* reporter assay, which includes the natural context of *GCN4* with all 4 uORFs (Figure 6B). Consistent with Ribo-seq and RNA-seq experiments (Figure 6A), the *GCN4-lacZ* reporter showed that *rad6*Δ cells have higher Gcn4 levels compared to WT cells (Figure 6B, left bar chart). As a positive control for our reporter, we treated the cells with 3-Amino-1,2,4-triazole (3-AT), which mimics amino acid starvation by inhibiting histidine (His) biosynthesis. 3-AT increases ribosome stalling at His codons and thereby activates Gcn2, leading to phosphorylation of eIF2α and higher Gcn4 translation (Klopotowski and Wiater, 1965). Expectedly, the *GCN4-lacZ* reporter showed increased expression upon 3-AT treatment in both WT and *rad6*Δ, but not *gcn2*11 cells (Figure S5A). As a separate control, we also included a construct with only uORF1, which exhibited increased expression of *GCN4* for both strains, due to the lack of uORF2-4 that would otherwise repress *GCN4* translation (Fig. 6B, middle bar chart, note change in axis). To test if the translation of *GCN4* is activated by leaky scanning in *rad6*Δ cells, we used another reporter where uORF1 is repositioned downstream and extended to overlap with the beginning of the *GCN4* main ORF. Under normal conditions, uORF1 should prevent scanning ribosomes from reaching the *GCN4* main ORF. The only way scanning ribosomes could reach the *GCN4 main* ORF would be if they scanned past uORF1 via “leaky scanning.” We did not see any difference in *GCN4-lacZ* expression from the uORF1-extended reporter between WT and *rad6*Δ cells (Figure 6B, right bar chart), suggesting that lack of Rad6 does not increase leaky scanning.

Mutations that cause *GCN4* translation to become constitutively active lead to a phenotype traditionally referred to as “Gcd^-^” (Hinnebusch, 2005). This phenotype, which describes the *rad6*Δ cells, could occur when TC assembly is impaired (lowering the availability of TC) or when the rate of TC re-loading onto 40S subunits after uORF1 is reduced. Lower overall translation level (Figures 4B and S3A) in *rad6*Δ cells in the absence of peroxide is indeed consistent with an interpretation where TC availability is low, which also results in constitutive translation of *GCN4*. One explanation for reduced TC availability would be lower expression of genes that maintain the TC. While we did not notice a change in the expression of these “Gcd” genes in our RNA-seq and Ribo-seq data (Figure S5B), other genes or other mechanisms (such as proteolysis) could mediate this effect.

Our data suggested this activation of *GCN4* translation in the absence of eIF2α-P may also be true in the presence of peroxide. We showed that *rad6*Δ cells that are peroxide-treated had a lower level of eIF2α-P compared with similarly treated WT cells (Figure 5C). However, the translational efficiency of *GCN4* was higher in these peroxide-treated *rad6*Δ cells (Figures 6A, pink bar higher than blue bar) and this was also true for the *GCN4-lacZ* reporter (S5C, gray bar with peroxide higher than black bar with peroxide). Moreover, RNA-seq data showed that known Gcn4-target mRNAs (Rawal et al., 2018) were upregulated at the transcriptional level in *rad6*Δ cells upon peroxide treatment compared with WT cells (Figure 6C). This indicates that the production of Gcn4 without eIF2α-P leads to the expected functional outcome. Our results therefore suggest a model where loss of ubiquitination by Rad6 causes dysregulation of eIF2α phosphorylation but that constitutive translation of *GCN4* in *rad6*Δ cells offers some compensation. The interplay of these effects may lead to altered fitness of *rad6*Δ cells during oxidative stress.

## DISCUSSION

Our work has revealed an oxidative stress response pathway that regulates translation through the E2 ubiquitin conjugase, Rad6. We found that Rad6 plays a key role in redox pausing of ribosomes as well as oxidative stress-induced translation inhibition and eIF2α phosphorylation, all contributing to the maintenance of cellular homeostasis during oxidative stress.

Cells induce complex regulatory programs in order to survive challenging conditions induced by oxidative stress. One such strategy is to inhibit global protein synthesis to direct cellular resources for the specific expression of survival genes. Although inhibition of translation initiation by eIF2α phosphorylation plays a key role in repressing translation during oxidative stress, previous data showed that even in the absence of Gcn2 (the sole eIF2α kinase in yeast), translation continues to be inhibited upon oxidative stress (Shenton et al., 2006). These data hinted that in addition to eIF2α-P, other pathways can also repress translation during and after initiation. We previously showed that Rad6-mediated K63-linked polyubiquitin chains change the conformation of the ribosome, and this conformation could block translation elongation (Zhou et al., 2020). Consistent with this idea, we found that oxidative stress induced by peroxide treatment leads to redox pausing at specific sequence motifs, particularly XIP, and this was dependent on Rad6 (Figures 1E-I and 2B-D). In addition, the redox pausing signatures were lost in the cells that express the catalytically dead Rad6 (Figures 2B-C), showing that it is the ubiquitination event that influences redox pausing. Strikingly, Rad6 was also necessary for a substantial portion of translational repression that is induced by oxidative stress (Figures 4B-C and S3), further supporting a potential role for redox pausing in the inhibition of protein synthesis. Although the mechanisms of redox pausing are not entirely clear, we ruled out changes to tRNA stability due to peroxide treatment (Thompson et al., 2008; Wu et al., 2019) by showing that the peroxide concentration used herein does not lead to Pro-tRNA^AGG^ degradation (Figure 4A). Altered modifications or the aminoacylation status of tRNA could possibly underlie redox pausing signatures. In particular, oxidative stress is known to affect tRNA modifications (Gu et al., 2014) and these changes in tRNA modifications influence ribosome pausing at the corresponding codons. For example, a yeast tRNA-Leu modification (m^5^C) was shown to be promoted by peroxide treatment, which leads to biased translation of genes containing Leu codons (Chan et al., 2012). Therefore, it is possible that increased or decreased modification of other tRNAs by peroxide could contribute to the P-site pausing signatures, such as Ile, observed in our data. These pausing events could be exacerbated when combined with a poor peptidyl-transfer substrate at the A-site, such as Pro. Interestingly, Pro acts as a scavenger of ROS (Ben Rejeb et al., 2014; Liang et al., 2013; Szabados and Savoure, 2010; Takagi, 2008) and therefore overall Pro amino acid levels (and tRNA aminoacylation) may also be affected by oxidative stress. In *S. pombe,* peroxide treatment was shown to cause ribosome pausing and collisions at Trp codons due to decreased levels of charged Trp-tRNA (Rubio et al., 2021). It is therefore possible that lower levels of charged Pro-tRNA, either due to diminished levels of Pro amino acids or an aminoacylation defect, could contribute to the A-site Pro redox pausing that we observed.

At least two models could explain how redox pausing events are eliminated in the absence of Rad6: (1) Ribosome ubiquitination by Rad6 alters a step in the elongation cycle. For example, it could change the rate-limiting step so that the time spent stalled on redox pausing motifs no longer makes up a significant part of the translation cycle. Loss of Rad6 would therefore lead to slower elongation under oxidative stress without any strong sequence-specific signature. However, this seems unlikely since the translation rate is faster, rather than slower, than expected (Figures 4B-C and S3) in *rad6*Δ cells under oxidative stress. Alternatively, Rad6-mediated ubiquitination could cause ribosome pausing and collisions during oxidative stress (i.e. at XIP motifs) and loss of Rad6 would eliminate these redox pausing events. Our findings support this second model, since translation is consistently faster in peroxide-treated *rad6*Δ cells compared to WT (Figures 4B-C and S3). (2) Rad6 could target transient redox pausing events (i.e. at XIP motifs) for ubiquitination, making them longer-lived and visible in our Ribo-seq data. We have shown that ribosomal ubiquitination during oxidative stress occurs independently of active translation (Back et al., 2019), supporting the fact that ribosomes paused at problematic sequences could be the preferred target of ubiquitination. Our study therefore establishes that Rad6-mediated ubiquitination affects elongation and offers models to guide future investigation of this phenomenon’s effect in the cell.

Even in the absence of oxidative stress, Rad6 is strongly associated with polysomes (Simoes et al., 2022), suggesting that Rad6 constitutively binds to ribosomes and ubiquitinates them, consistent with the notion that redox inhibition of the deubiquitinating enzyme Ubp2 seems to drive ubiquitin chain accumulation (Silva et al., 2015). The fate of these ubiquitinated ribosomes, however, remains unclear. One possibility is that they are rescued, perhaps by proteins in the RQC pathway. However, our rescue-based reporter experiments showed that lack of Rad6 did not affect the Fluc/Rluc ratio in the reporter containing the redox pausing motif XIP (Figure 2A), suggesting that ubiquitination of ribosomes by Rad6 does not lead to rescue. This finding is also consistent with the previous results showing that K63-ubiquitinated ribosomes are detected in polysomes and are therefore engaged in translation (Silva et al., 2015). The other, and more plausible scenario is that ubiquitin marks are removed from the ribosome once the oxidative stress insult is no longer present. We favor this possibility since the deubiquitinase, Ubp2, removes K63 ubiquitin on the ribosome and the activity of this enzyme is mediated by peroxide levels in the cell (Silva et al., 2015).

Although lack of Hel2 did not affect redox pausing signatures and K63-linked ubiquitination (Figures 3A-B), it is possible that oxidative stress induced by other ROS could lead to ribosome collisions that are targeted by RQC. For example, it was shown that the treatment of cells with 4-nitroquinoline 1-oxide (4-NQO), which is proposed to produce superoxide and hydroxyl radicals after enzymatic process (Arima et al., 2006), increases oxidation of guanines and leads to ribosome stalling that is detected by the RQC pathway (Yan et al., 2019). Therefore, different ROS, their modes of production, abundance, and subcellular location could potentially engage unique pathways of translation control mediated by Hel2 and Rad6 pathways.

In addition to RQC, we also examined the relationship of Rad6 with the ISR. Peroxide was known to trigger the ISR and we and others have shown that the ISR can also be activated by ribosome collisions in yeast (Meydan and Guydosh, 2020; Yan and Zaher, 2021), which induces eIF2α phosphorylation. Our data in *rad6*Δ cells show that the ISR pathway is functional and can be activated by starvation of histidine (Figure S5A). Our observation that Rad6-mediated stalling is induced by peroxide treatment therefore offers an explanation for how oxidative stress leads to eIF2α phosphorylation. The loss of redox pausing in the absence of Rad6 may account for the lack of eIF2α phosphorylation. In combination with the loss of redox pausing, lower levels of oxidative stress-induced eIF2α-P could also help explain the lack of translational repression in peroxide-treated *rad6*Δ cells (Figures 4B-C and S3). Interestingly, although the effect of Rad6 on redox pausing was reproducible in both SUB280 and S288C strains, we observed impaired induction of eIF2α phosphorylation by peroxide only in the SUB280 strain (Figure S4E). This suggests that other cellular inputs contribute to eIF2α phosphorylation in addition to Rad6-mediated redox pausing and these other inputs could be more dominant in the S288C strain. Consistent with this idea that ISR regulation may vary somewhat between strains, it has been reported that Gcn4 translation is not activated by peroxide in the S288C strain (Picazo and Molin, 2021). In addition, S288C has a unique genetic background that affects mitochondria physiology and cellular redox biology (Young and Court, 2008), which could influence redox experiments in this strain. Despite impaired eIF2α phosphorylation, Gcn4 is still constitutively translated (Figure 6) in *rad6*Δ cells (Gcd^-^ phenotype) even in the absence of oxidative stress, which is potentially indicative of an initiation defect (Hinnebusch, 2005). An initiation defect would also be consistent with the slower growth of *rad6*Δ cells (Simoes et al., 2022) and overall lower translation rate in the absence of peroxide (Figures 4B-C and S3). One possibility for the Gcd^-^ phenotype could be that the lack of Rad6 affects the expression of “Gcd” genes and decreases their availability, resulting in uORF bypassing and increased translation of *GCN4*. However, we found that the expression of these genes was not affected in *rad6*Δ cells (Figure S5B) and therefore other mechanisms drive the constitutive translation of *GCN4*. We previously observed increased presence of subunits of the translation initiation factor eIF3 associated with the WT compared to cells unable to produce K63-linked ubiquitin chains (Back et al., 2019). It is therefore possible that ubiquitination of translation factors by Rad6 may regulate initiation. Finally, the lack of Rad6 could lead to increased *GCN4* translation by promoting reinitiation over recycling at uORF1. Although the exact mechanisms by which Rad6 affect *GCN4* translation are yet to be identified, the potential role of Rad6 in translation initiation expands its functions beyond translation elongation. It remains unclear whether translational control at initiation and elongation mediated by Rad6 occurs independently and whether both contribute to cellular fitness during oxidative stress.

Rad6 is conserved from yeast to human cells and mutations in its human homolog, UBE2A, are linked to intellectual disability type Nascimento due to loss of UBE2A activity (Czeschik et al., 2013; Nascimento et al., 2006). UBE2A was also shown to modulate neuronal function in flies by interacting with the E3 ubiquitin ligase Parkin and thereby inducing mitophagy (Haddad et al., 2013). We previously showed that Rad6 carrying the corresponding disease mutations in its human homolog leads to dysregulated K63 ubiquitination response during oxidative stress in yeast (Simoes et al., 2022), further supporting the idea that the role of Rad6 in cellular homeostasis is conserved across species. Our studies in yeast, therefore, reveal crucial insights into the cellular response to UBE2A deficiency and could be important for delineating the disease mechanisms.

## AUTHOR CONTRIBUTIONS

Methodology, Validation, Visualization, S.M., G.C.B., V.S.; Resources, L.H., B.K.C.; Software and Formal analysis, S.M and N.R.G; Conceptualization, Writing, S.M., G.C.B., V.S, N.R.G and G.M.S.; Supervision, Project Administration, and Funding Administration, N.R.G and G.M.S.

## DECLARATIONS OF INTEREST

The authors declare no competing interests.

## ACKNOWLEDGEMENTS

We thank Yan Luo, Yuesheng Li, Patrick Burr (NIH/NHLBI DNA Sequencing and Genomics Core), Ilhan Akan and Harold Smith (NIH/NIDDK Genomics Core) for sequencing experiments. We thank David Young and Agnes Karasik for help with the northern blot experiments and Ayse Ecer for help with analysis of publicly available oxidative stress profiling datasets. We thank Alan Hinnebusch for kindly providing *GCN4* reporter plasmids and Thomas Dever for providing anti-eIF2α antibodies. We thank Xinnian Dong, John Panepinto, Corey Knowles, Lucia Strader, and Marcelo Figueiredo for making resources and equipment available. We thank the Flow Cytometry core in the Duke Cancer Institute. We thank Alan Hinnebusch, Jon Lorsch, Thomas Dever, NIDDK Laboratory of Biochemistry and Genetics. We also thank the members of the Guydosh and Silva Labs for feedback on our project. The work is supported by the Intramural Research Program of the NIH, National Institute of Diabetes and Digestive and Kidney Diseases (NIDDK; DK075132 to N.R.G.), the Postdoctoral Research Associate Training Program (PRAT) at the National Institute of General Medical Sciences (NIGMS) (1FI2GM137845 to S.M.) and US National Institutes of Health R35 Award GM137954 (G.M.S.).

## EXPERIMENTAL MODEL AND SUBJECT DETAILS

The yeast strains used in this study are listed in Table S1. SUB280 strain derivatives were grown in synthetic defined (SD) medium composed of D-Glucose (BD Difco, #215510), yeast nitrogen base (BD Difco, #291940) and drop-out amino acid medium without Leu and Trp (Sigma, #Y0750). SUB280 *rad6*Δ *RAD6* and SUB280 *rad6*Δ *RAD6(C88A)* cells were grown in SD media supplemented with drop-out amino acid supplements without Leu, Trp and Ura (Sigma, #Y1771). S288C strain derivatives were grown in SD complete media by using drop-out amino acid supplements without Leu and Trp (Sigma, #Y0750) and supplementing it back with L-leucine (Sigma, #L800) and Tryptophan (Sigma, #T8941). Starter cultures were grown at 30°C overnight and then diluted to an OD_600_ of 0.001 (*rad6*Δ cells) or 0.0001 (WT cells) and were grown to a final OD_600_ between 0.5 and 0.6 for ∼16 hours. Unless noted otherwise, the cultures were treated with freshly diluted H_2_O_2_ (peroxide) (Sigma, #216763), achieving a final concentration of 0.6 mM, for 30 minutes, filtered and frozen in liquid nitrogen for Ribo-seq, Disome-seq and RNA-seq experiments.

## METHOD DETAILS

### Ribo-seq, Disome-seq and RNA-seq experiments

Ribo-seq, Disome-seq and RNA-seq experiments were performed based on published protocols (Guydosh and Green, 2014; McGlincy and Ingolia, 2017; Meydan and Guydosh, 2020). Frozen yeast cell pellets and frozen droplets of lysis buffer (20 mM Tris pH 8.0, 140 mM KCl, 1.5 mM MgCl2, 1% Triton X-100 and 0.1 mg/mL cycloheximide [Sigma, #C7698]) were lysed using a Retsch Cryomill (Retsch 20.749.0001). The resulting powder of frozen cell and lysis buffer mixture was thawed at room temperature, transferred to a 50 mL falcon tube to spin at 3000 g for 5 min at 4°C. The supernatant was then spun at 21000 g for 10 min at 4°C. The absorbance of the supernatant (cell lysate) at 260 nm was recorded and total “OD” of the lysate was calculated as the product of the volume (in mL) multipled with A260 reading. A fraction of the lysate equivalent to OD=45 was flash frozen in liquid nitrogen. Prior to RNase I digestion, lysates were thawed, diluted with an equal volume of lysis buffer and then digested with ∼26 U of RNase I (Ambion, #AM2294) per OD for 1 h at room temperature (22°C) with gentle agitation at 700 rpm. Monosome (for Ribo-seq) and disome (for Disome-seq) fractions were separated by loading the lysates onto a 10%-50% sucrose gradient, prepared in gradient buffer (final concentration: 20 mM Tris pH 8.0, 150 mM KCl, 5 mM MgCl2, 0.5 mM DTT), and spun at 40,000 rpm for 3 hours at 4°C using a SW 41 Ti Swinging-Bucket Rotor (Beckman Coulter). Sucrose gradient fractionation was performed by using a Brandel Density Gradient Fractionation System. The peaks corresponding to monosomes and disomes were collected and RNA was purified by using the SDS, hot acid phenol-chloroform extraction method. For RNA-seq, the total RNA was isolated directly from the frozen cell pellets by the SDS, hot acid phenol-chloroform extraction method and fragmented in a buffer (pH 9.2) containing 12 mM Na2CO3, 88 mM NAHCO3, 2 mM EDTA for 35 minutes at 95°C. The total RNA was cleaned up using the Oligo Clean & Concentrator kit (Zymo Research, #D4060). Monosome/disome footprints and total RNA isolated as described above were run on a 15% TBE-Urea polyacrylamide gel (Bio-Rad, #3450091) for the size selection process. For Ribo-seq, Disome-seq and RNA-seq, RNA fragments between 25-34 nt, 54-68 nt and 50-70 nt were excised from the gel, respectively. We used the 50 nt band from a small RNA marker (Abnova, #R0007) for RNA-seq experiments and other RNA size markers used for size selection are listed in Table S2. The excised gel pieces were frozen on dry ice for 30 minutes and thawed in RNA extraction buffer (0.3 M NaOAc, 1 mM EDTA, 0.25% SDS) overnight at 20°C with gentle agitation (700 rpm). Next day, RNA was precipitated and the pellet was resuspended in 10 mM Tris pH 8.

### Next generation sequencing library preparation

Library preparation was conducted by following published protocol (McGlincy and Ingolia, 2017). The RNA fragments from Ribo-seq, Disome-seq and RNA-seq experiments were first dephosphorylated using PNK (NEB, #M0201L) and ligated to preadenylated linkers containing a 5 nt-long random Unique Molecular Index (‘UMI’) and a 5 nt barcode that is unique for each sample (listed in Table S2). The linkers that were pre-adenylated using a 5’ DNA adenylation mix (NEB, #E2610L) were ligated to dephosphorylated RNAs using T4 truncated RNA ligase 2 (K227Q) (NEB, #M0351L). Unligated linkers were depleted by using 5 U per sample of 5’ deadenylase (NEB, #M0331S) and RecJ exonuclease (Biosearch Technologies, #RJ411250). Ligated RNA samples with unique barcodes were pooled and cleaned up using the Oligo Clean & Concentrator kit (Zymo Research, #D4060). All samples were next reverse transcribed by using Superscript III (Invitrogen; 18080044), and the reverse transcription primer (NI-802, listed in Table S2) containing a random 2 nt UMI. At this step, ribosomal RNA (rRNA) was removed from Disome-seq and RNA-seq samples by using Qiagen FastSelect (Qiagen, #334215). The cDNAs obtained from this reaction were resolved on a 10% TBE-Urea gel (Bio-Rad, #3450089) and cDNAs were extracted using DNA gel extraction buffer (0.3 M NaCl, 1 mM EDTA, 10 mM Tris pH 8) with gentle agitation (700 rpm) overnight at 20°C. The next day, DNA was precipitated and the pellet was resuspended in 10 mM Tris pH 8. The footprints were circularized using CircLigase ssDNA Ligase (Biosearch Technologies, #CL4115K). For Ribo-seq samples, rRNA removal was performed at this stage by oligonucleotide substraction using Dynabeads MyOne Streptavidin C1 (Invitrogen, #65001) and DNA oligos that are the reverse complement of ribosomal RNAs (listed in Table S2). The samples were then amplified by PCR using Phusion DNA Polymerase (ThermoFisher Scientific, #F530L) and resulting product were resolved in hand-poured 8% native TBE gel. The libraries were extracted using DNA gel extraction buffer (0.3 M NaCl, 1 mM EDTA, 10 mM Tris pH 8) with gentle agitation (700 rpm) overnight at 20°C. The next day, DNA was precipitated and the pellet was resuspended in 10 mM Tris pH 8 to obtain the final library. For Disome-seq of *rad6*Δ cells, four different PCR libraries were pooled to increase the yield due to lower levels of disome population in these cells. Quality of the library was assessed by using a BioAnalyzer via the High Sensitivity DNA Kit (Agilent, #5067-4626) and TapeStation via High Sensitivity D100 Screen Tape System (Agilent, #5067-5584, #5067-5585). Sequencing experiments were performed by the NIDDK Genomics Core and NHLBI DNA Sequencing and Genomics Core at NIH (Bethesda, MD). Sequencing of SM099F, SM100F, SM103F-SM110F samples was conducted on an Illumina HiSeq2500 machine (single end, 50 bp cycle) and the rest of the samples on an Illumina NovaSeq machine (single end, 100 bp cycle).

### Computational processing and analysis of Ribo-seq, Disome-seq and RNA-seq data

The sequencing data was processed as described previously (Meydan and Guydosh, 2020). Custom scripts are available on Github (https://github.com/guydoshlab). Briefly, fastq files of sequencing samples were provided by NIDDK Genomics Core and NHLBI DNA Sequencing and Genomics Core (NIH). We used Cutadapt (Martin, 2011) to remove linkers and demultiplex for retrieving individual samples from pooled data. For RNA-seq samples, we trimmed all the reads to 50 nt by using following parameters: -j 0 - l 57 --discard-untrimmed. To remove rRNA and tRNA reads, we then aligned the files to an index of noncoding RNAs with Bowtie version 1.1.2 (Langmead et al., 2009) by using following parameters: -v 2 -y -S -p 12. We removed PCR duplicates by using a custom python script. We then aligned the deduplicated files to coding regions and splice junctions of R64-1-1 S288C reference genome assembly (SacCer3, Saccharomyces Genome Database Project) by using the following parameters: -v 1 -y -a -m 1 --best -- strata -S -p 4. The number of reads that were obtained after each of these steps are outlined in Table 3.

Custom python scripts are used for the data analysis by using biopython version 1.72 and python 2.7.18. For Ribo-seq and Disome-seq experiments, only the reads between 25-34 and 57-63 nt were analyzed, respectively. Ribo-seq and Disome-seq reads were aligned by their 3’ ends. For RNA-seq experiments, 50 nt reads were analyzed and coverage of reads was used instead of 3’ alignment. All reads were normalized in units of rpm (reads per million mapped reads), which was computed by dividing the read count at each nt position by the total number of mapped reads and then multiplying the result with 10^6^.

Quantitation of Ribo-seq and RNA-seq data was performed by summing the total number of normalized reads mapping to each coding sequence or UTR regions obtained from published studies (Ng et al., 2020; Pelechano et al., 2014). These total number of reads per gene was normalized by the gene’s length (in kilobases) to obtain rpkm values. Ribo-seq reads were shifted 15 nt from their 3’ end to align the P-site to the beginning of each gene. Data from 15 nt of either end of the ORFs was eliminated to reduce the effects of initiation and termination on ribosome occupancy. For differential expression analysis by DESeq2 (Love et al., 2014), raw counts were first generated for each gene. The gene expression profiles were compared by running DESeq2 on Rstudio and Padj values were obtained. We used Padj < 0.05 for significance cut-off, log_2_FoldChange value > 0.8 for upregulated and < -0.8 for the downregulated genes. Volcano plots were generated by using ggplot2 in Rstudio (Wickham, 2016). Gene ontology analysis was performed by using PANTHER Classification System (http://www.pantherdb.org/) (Mi et al., 2013; Thomas et al., 2022) with following parameters: PANTHER version 17.0 Overrepresentation Test, FISHER test with FDR correction, PANTHER GO-Slim Biological Process with Saccharomyces Cerevisiae - REFLIST (6050) as a reference gene list. Gcn4-target mRNAs were obtained from a published ChIP-seq dataset (Rawal et al., 2018). From this dataset, the first 250 genes that had > 2 fold increase in Rbp3 (RNA polymerase B) occupancy in starved cells with reproducible induction by Gcn4 in other datasets (Natarajan et al., 2001; Qiu et al., 2016; Saint et al., 2014) were defined as Gcn4 targets.

Metagene plots were generated by averaging rpm around the start and stop codons normalized by the total number of reads in a given window for each gene (100 nt upstream of the ORF and 300 nt into the ORF for start codon metagene; 300 nt of the ORF and 100 nt downstream of the ORF for stop codon metagene). ORFs that were unidirectionally overlapping with other ORFs, the genes with features smaller than the window size, and the genes without any mapped reads were excluded from the analysis. Average reads plots of XIP motifs were generated by first creating a list of occurrences of XIP motifs in the yeast transcriptome and then averaging normalized monosome or disome occupancy from a region of interest (50 nt upstream and 50 nt downstream of XIP motif). Normalization was done by dividing the rpm at each position in the region of interest by the average rpm of the gene.

Pause scores were computed by dividing the rpm of a motif by the average rpm in a region of interest (±50 nt of each motif). Pause scores for sites that are smaller than the ±50 nt window were eliminated from the analysis. Average pause scores were generated by averaging the individual pause scores for each tri-amino acid motif. We excluded the motifs that were represented in the genome less than 100 times to reduce noise, which resulted in 6267 motifs that were compared across datasets. Individual pause scores for XIP motifs were visualized in a box plot to show the distribution and significance of XIP pause scores. The significance of differences in the median of these individual pause scores were computed by independent 2-group Mann Whitney U Test in Rstudio.

### Dual luciferase reporter experiments

#### 1. Plasmids Building

RLuc-P2A-X-P2A-Fluc plasmids (where X represents a variable sequence) were assembled using NEB Builder HiFi DNA Assembly Cloning Kit (New England Biolabs, #E2621S) by combining the original plasmid digested with HindIII (New England Biolabs, #R3104S) and NotI (New England Biolabs, #R3189S) and the gene fragments and oligo listed in Table S2.

#### 2. Luminescence activity measurement

Yeast strains transformed with the plasmids described above in log phase grown in SD-Ura medium were pelleted down and transferred to SD-Ura-Met to induce plasmid expression for 90 minutes. For cells treated with indicated H_2_O_2_ concentrations, plasmid expression was induced for 60 minutes and then H_2_O_2_ was added to the medium and incubated for 30 minutes under agitation. Pelleted cells were disrupted by glass beads agitation at 4°C in 1x Passive Lysis Buffer provided in the Dual-Luciferase® Reporter Assay System (Promega, #E1910). Extracts were clarified by centrifugation, and protein concentration was determined by BCA assay (ThermoFisher, #23225). The luminescence activities of RLuc and FLuc were collected for 5 µg of protein mixed with the respective substrates. For Figures 3C and S2C, luminescence values were obtained in a VictorX (PerkinElmer) plate reader. For Figure S1C luminescence values were obtained using a Glo Max® (Promega) plate reader. For Figures 2A and S1D luminescence values were obtained using a CLARIOstar Plus (BMG LabTech) plate reader.

### Northern blotting

The tRNA-Pro coding sequence was ordered as gBlock (listed in Table S2) and was assembled with digested YCplac33 backbone using NEBuilder HiFi DNA Assembly Cloning Kit (NEB, #E5520). The tRNA-Pro probe sequence was amplified from this plasmid using the primers in Table S2 and in vitro transcribed by using Digoxigenin-11-UTP included in DIG Northern Starter Kit (Sigma, #12039672910). 25 µg total RNA each from WT and *rad6*Δ cells was resolved on 15% TBE-Urea polyacrylamide gel (Bio-Rad, #3450091). The RNAs were then transferred onto positively charged nylone membrane (Sigma, #11209299001) in 20x SSC buffer for 3 hours by using Nytran SuPerCharge turboblotter system (Cytiva, #10416302) following manufacturer’s instructions. The RNA was UV-crosslinked to the membrane by using VWR UV crosslinker (VWR, #89131-484) at 120,000 microjoules per cm^2^ and 100 ng/mL of the probe diluted in DIG Easy Hyb Granules Working Buffer was hybridized overnight at 42°C with gentle agitation (Sigma, #11796895001). Next day, the membranes were washed with low stringency wash buffer (2X SSC, 0.1% SDS) and then with high stringency wash buffer (1X SSC, 0.1% SDS), twice for 5 minutes at room temperature for each. The membrane was washed and subjected to Anti-DIG-AP antibody by using DIG Wash and Block Buffer Set (Sigma, #11585762001) and immonological detection of the membrane was conducted by CDP-Star chemiluminescent substrate included in the northern blotting kit.

### Translation rate assays

Indicated yeast strains in logarithmic phase grown into SD medium were back-diluted to OD_600_ 0.1-0.2 in SD-Met medium. At OD_600_ 0.4-0.5, cells were treated with 50 µM of HPG (L-Homopropargylglycine, Sigma, #900893) and collected by centrifugation after 15, 30, 45, and 60 minutes of incubation at 30°C under agitation. For H_2_O_2_ treatment, cells were incubated with 0.6 mM of H_2_O_2_ for 15 minutes prior to HPG incubation as above. Pelleted cells were fixed overnight in 70% ethanol at 4°C and the HPG conjugation with Alexa Fluor 488 was done using the Click-iT® HPG Alexa Fluor® Protein Synthesis Assay (ThermoFisher, #C10428) following manufacturer’s instructions. Alexa Fluor 488 fluorescent signal was measured in the BD FACS Canto flow cytometer using a 488 nm laser. Single-cell population gates, histograms plots, and mean/median calculations were done using FlowJo software (Becton Dickinson).

### Western blotting

For blot in Figures 3B and 5C-E, yeast cells grown to logarithmic phase (OD∼0.5-0.6) were disrupted by glass-bead agitation at 4°C in buffer containing 50 mM Tris-HCl pH 7.5, 150 mM NaCl, 20 mM iodoacetamide, 1X protease inhibitor cocktail set I (Sigma, #539131). Extracts were clarified by centrifugation, and protein concentration was determined by Bradford assay (Bio-Rad, #5000205) prior to western blotting. Proteins were separated by standard 10% or 12.5% SDS-PAGE loaded in Laemmli buffer and transferred to PVDF membrane (ThermoFisher, #88518). Immunoblotting was performed using the following antibodies: anti-K63 ubiquitin (EMD Millipore, #051308), anti-GAPDH (Abcam, #ab9485), anti-eIF2α-phospho (Cell Signaling, #3398), anti-actin (Cell Signaling, #4967). For the blot in Figure S4E, yeast extracts were prepared from 25 mL of yeast cells grown to logarithmic phase (OD∼0.5-0.6) by TCA precipitation. 10 µL samples were loaded on 4-20% Mini-Protean TGX gel (Bio-Rad, #4561096) and transferred to a PVDF membrane (Bio-Rad, #1704156). The proteins were detected using antibodies against eIF2α-phospho (Abcam, #32157). The antibody against yeast eIF2α was kindly provided by the laboratory of Thomas Dever (NIH/NICHD).

#### *GCN4-lacZ* reporter assays

Expression of *GCN4-lacZ* fusions was measured by assaying β-galactosidase in whole-cell extracts. Yeast cells transformed with GCN4-lacZ plasmids were grown to logarithmic phase (OD∼0.4-0.5) and disrupted by glass-bead agitation at 4°C in buffer containing 1x PBS, 40 mM KCl, and 10 mM MgCl2. Extracts were clarified by centrifugation, and protein concentration was determined by BCA assay (ThermoFisher, #23225). 120 µg protein was mixed with substrate containing 15mM ONPG (2-Nitrophenyl-β-D-galactopyranoside, Goldbio, #N27510), 5mM DTT, 1x PBS, 40mM KCl, and 10mM MgCl2, and incubated for 30 minutes at 37°C. Absorbance was read at 420 nm in a Tecan Sunrise plate reader. When noted, cells were treated with 0.6 mM H_2_O_2_ for 2 hours or 30 mM 3-amino-triazole (3-AT) for 5 hours in SD media without histidine.

---

**Figure S1.**
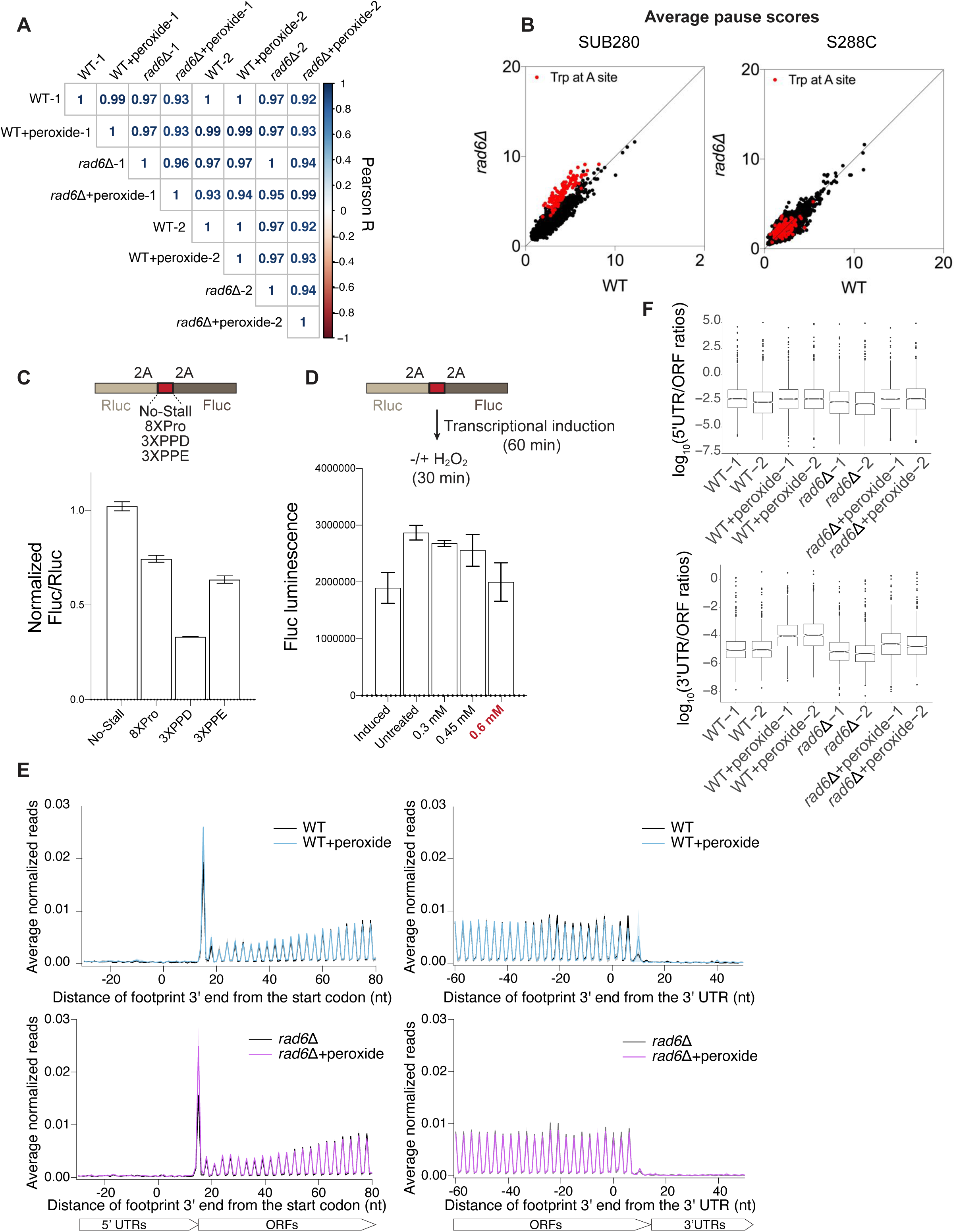
Global analysis of translation in WT and *rad6*ll. cells. Related to Figures 1 and 2. **(A)** Correlation matrix of the RNA-seq data corresponding to 2 replicates of WT and *rad6*Δ ± H_2_O_2_ (peroxide), computed using unnormalized counts for each gene. This shows the reproducibility of replicates and extent of change between different conditions. **(B)** Average pause scores of 6267 tripeptide motifs plotted for untreated WT and *rad6*Δ cells in either SUB280 (left) or S288C (right) background. Motifs with Trp at A-site are indicated in red. Note that Trp stalling in *rad6*Δ cells is specific to SUB280 strain. **(C)** Schematic of reporter experiments that test motifs with high pause scores in Ribo-seq data (top). Fluc/Rluc values of the reporters with stall-inducing sequences relative to the reporter without any stall sequence (No-Stall) shows that stalling motifs cause 20-70% reduction in Fluc, indicative of ribosome rescue or drop-off prior to Fluc (bottom). **(D)** Schematic of reporter experiments in the presence of peroxide (top). Transcription is induced for 60 min and then the cells are treated with peroxide for 30 min. The data (bottom) show firefly luminescence (only) in untreated cells and cells treated with 0.3, 0.45 and 0.6 mM of peroxide. This value increases during the induction phase (induced vs untreated bars) as the cells move toward a steady state. The dynamics of this process make it difficult to assess additional changes due to peroxide, which appears to reduce Firefly luminescence, and this effect is most severe for the concentration of interest (0.6 mM, indicated in red). **(E)** Metagene analysis showing the average normalized Ribo-seq reads mapped to genes that were aligned by their respective start codon (left panels) or stop codon (right panels) of WT (top) and *rad6*Δ (bottom) cells ± peroxide. Average of two replicates ± standard deviation (shaded) is plotted. Loss of Rad6 does not change the overall translation trends. **(F)** Box plots showing the distribution of 5’UTR/ORF (top) or 3’UTR/ORF (bottom) ratios for WT and *rad6*Δ cells ± peroxide. 5’UTR and 3’UTR translation slightly increases with peroxide treatment, and these trends are mostly similar in *rad6*Δ cells.

**Figure S2.**
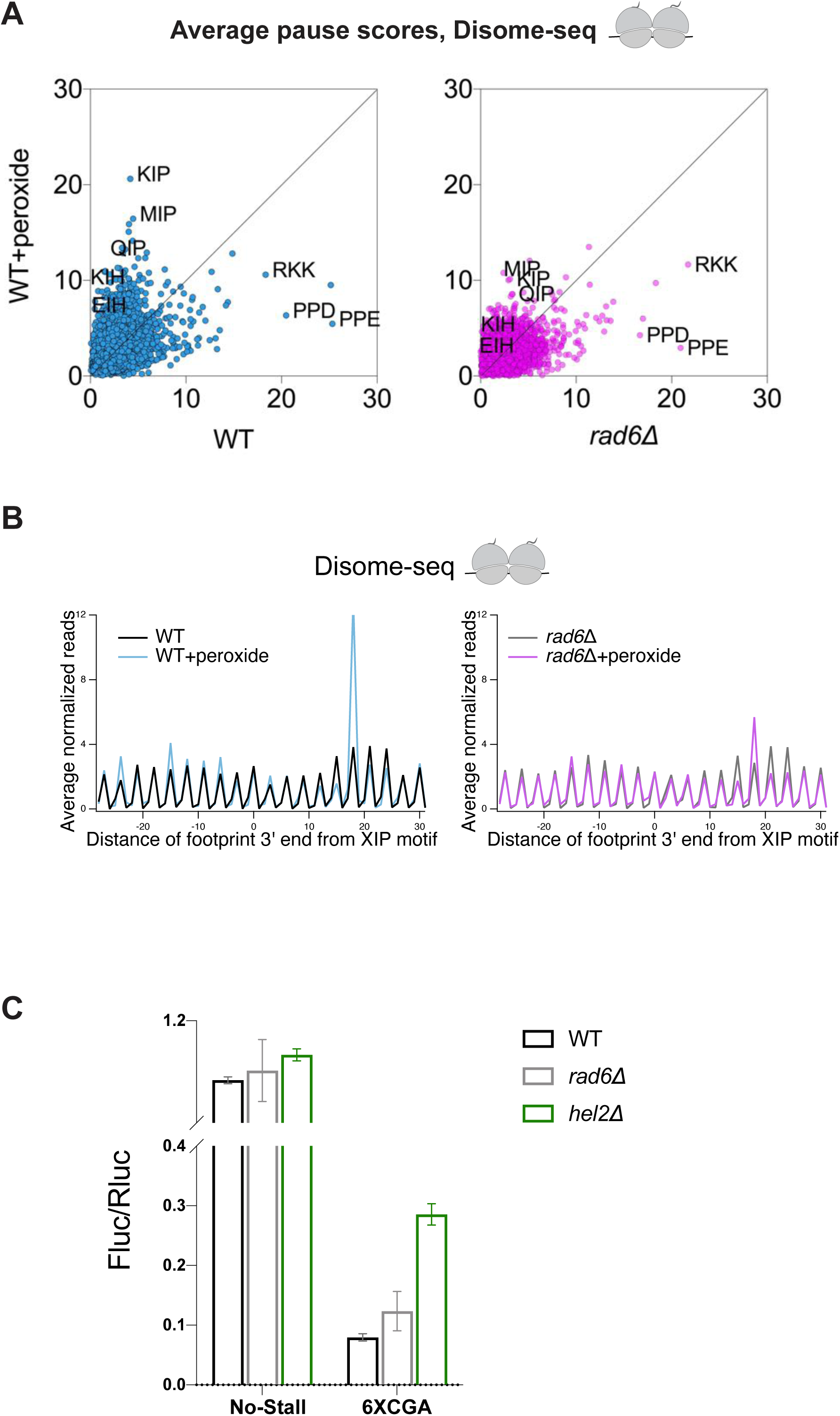
Disome-seq signatures of WT and *rad6*ll. cells and detailed RQC reporter results. Related to Figures 1 and 3. **(A)** Average Disome-seq pause scores of 6267 tripeptide motifs plotted for WT and *rad6*Δ cells ± peroxide. The redox pausing signatures are similar to Ribo-seq motifs. Average of two replicates is plotted. Motifs with Trp codons are excluded from this graph. **(B)** Average normalized Disome-seq reads mapped to genes aligned by their respective XIP motifs in WT (left) or *rad6*Δ (right) cells. Note that the XIP pausing by disomes is diminished in *rad6*Δ cells, consistent with results for Ribo-seq experiments. **(C)** Fluc/Rluc ratio of the dual luciferase reporter with 6XCGA shows that deletion of *HEL2* increases bypassing of the 6XCGA sequence, as anticipated. Loss of the *RAD6* gene, in contrast, does not affect this reporter.

**Figure S3.**
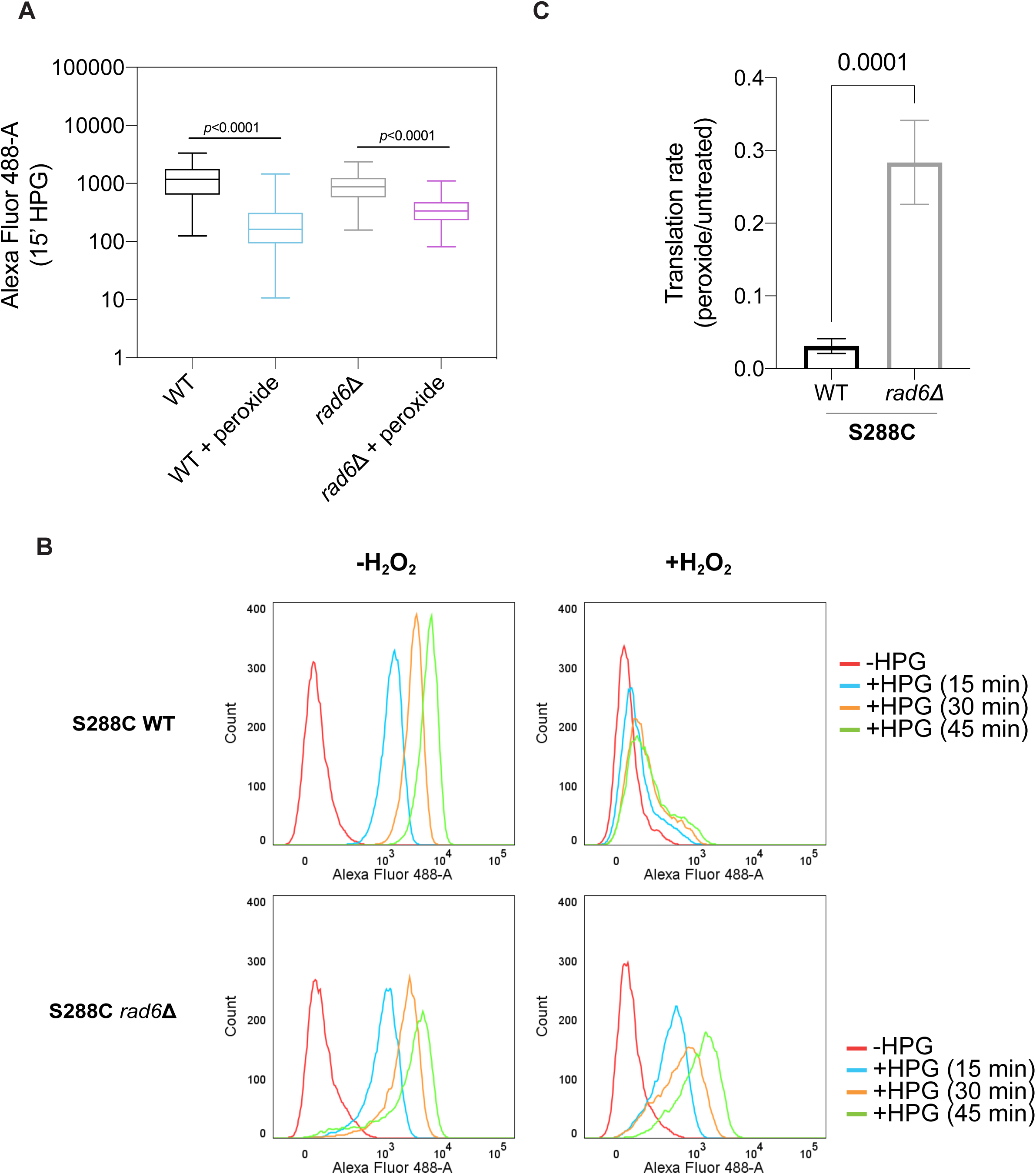
Translation rate assays in WT and *rad6*ll. cells. Related to Figure 4. **(A)** Raw fluorescence data for HPG incorporation at 15 minutes show that peroxide-induced translation inhibition is lower in *rad6*Δ cells compared to WT. The statistical significance is assessed by comparing the means via two-way Anova test. **(B)** Histograms of the HPG incorporation assay showing the number of cells (y-axis) and fluorescence measurements (x-axis) at indicated time points of HPG incubation (15-45 minutes). S288C WT cells exhibit decreased HPG incorporation in the presence of peroxide (top panel) and this inhibition is slowly released over time. In contrast, HPG incorporation in *rad6*Δ cells is affected less by the peroxide treatment (bottom panel). These data show the responses observed in the SUB280 strain are consistent in the S288C strain. **(C)** Quantification of HPG incorporation during peroxide treatment is shown as a normalized rate for HPG incorporation in treated vs untreated S288C cells. The translation rates were calculated by fitting the mean fluorescence values to a linear regression as a function of time. Significance is determined by two-tailed unpaired t-test. These data show the responses observed in the SUB280 strain are consistent in the S288C strain.

**Figure S4.**
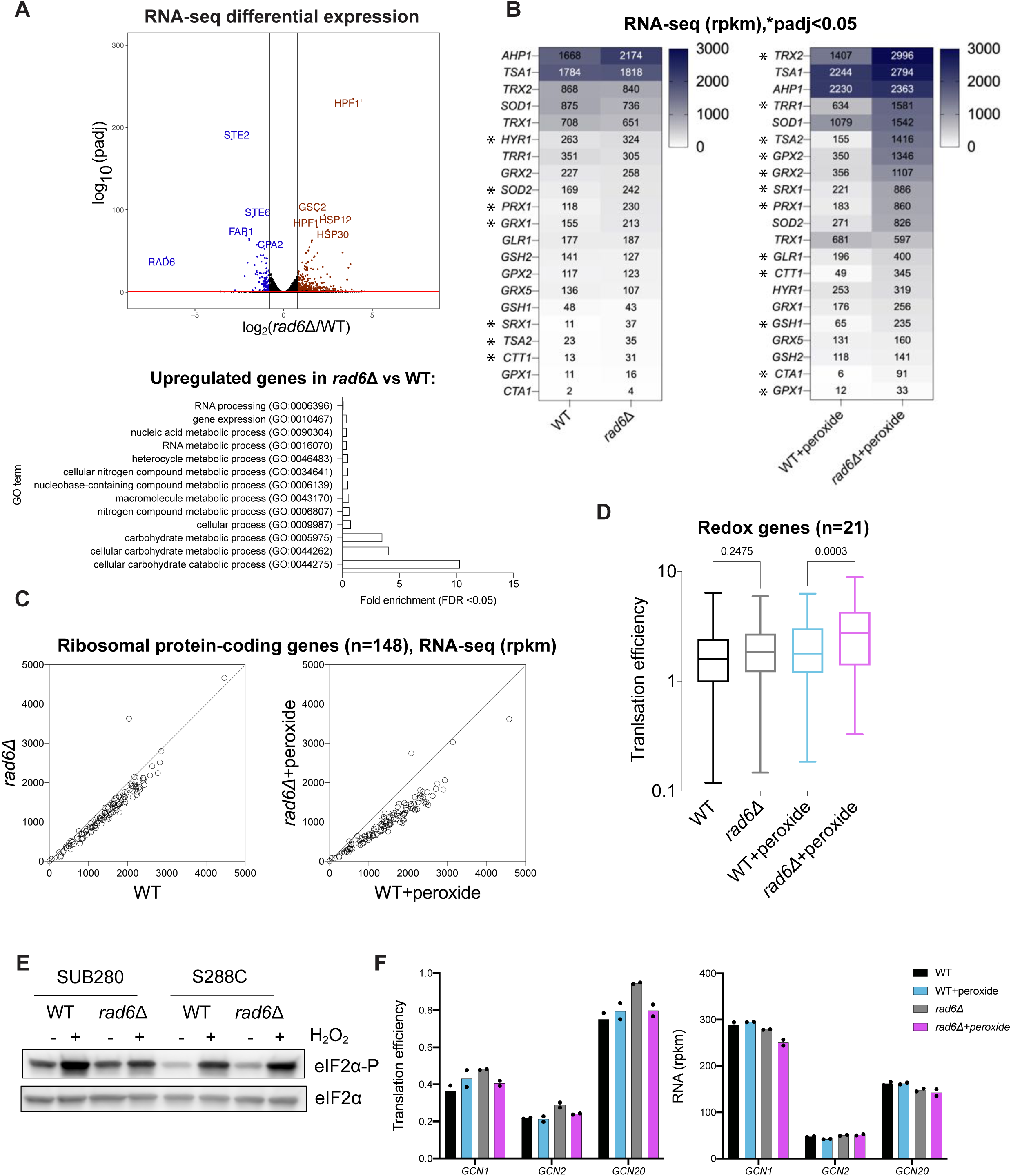
Differential expression analysis of WT and *rad6*ll. cells. Related to Figure 5. **(A)** Volcano plot showing the differential RNA expression in *rad6*Δ versus WT cells. Genes that are significantly upregulated (log_2_Fold Change > 0.8, padj < 0.05) or downregulated (log_2_Fold Change < -0.8, padj < 0.05), as determined by DESeq2 analysis are shown in red and blue, respectively. The significance cut-off is indicated with a red bar. Significantly enriched gene ontology (GO) terms of the genes that are significantly upregulated in the absence of Rad6 are shown at the bottom. GO analysis is conducted in PANTHER, GO-Slim Biological Process by using *Saccharomyces cerevisiae* (all genes in database) as reference list and test type as FISHER with FDR correction. Genes that are significantly downregulated in *rad6*Δ cells did not have a significantly enriched GO term. **(B)** Heatmap showing the RNA-seq expression (rpkm) of redox genes in the WT and *rad6*Δ ± peroxide samples. The plotted data shows the average of two RNA-seq replicates. The genes that are differentially expressed (DESeq2, padj < 0.05) are indicated with an asterisk (*). **(C)** RNA-seq data of ribosomal protein-encoding genes (n=148, gene names obtained from SGD) in WT vs *rad6*Δ cells ± peroxide show that mRNAs that encode ribosomal proteins are lower in *rad6*Δ cells, which is a hallmark of TOR pathway inactivation. **(D)** Translation efficiency (Ribo-seq reads normalized by RNA-seq reads) of 21 mRNAs that encode redox enzymes shows translational upregulation of these genes in *rad6*Δ vs WT cells in the presence of peroxide. Significance is calculated by one-way Anova test. ns = not significant. **(E)** Western blotting demonstrates that the phosphorylation of eIF2α due to peroxide is minimal in *rad6*Δ cells in the SUB280 background but not in the S288C background, suggesting that other inputs regulate eIF2α phosphorylation in the S288C background. Note that total eIF2α levels do not change with peroxide and in the different strains. **(F)** Bar graphs of the translation efficiency (left) and RNA levels (right) of *GCN1, GCN2*, *GCN20* genes show that the abundance and translation level of the mRNAs encoding these proteins are not affected by loss of Rad6.

**Figure S5.**
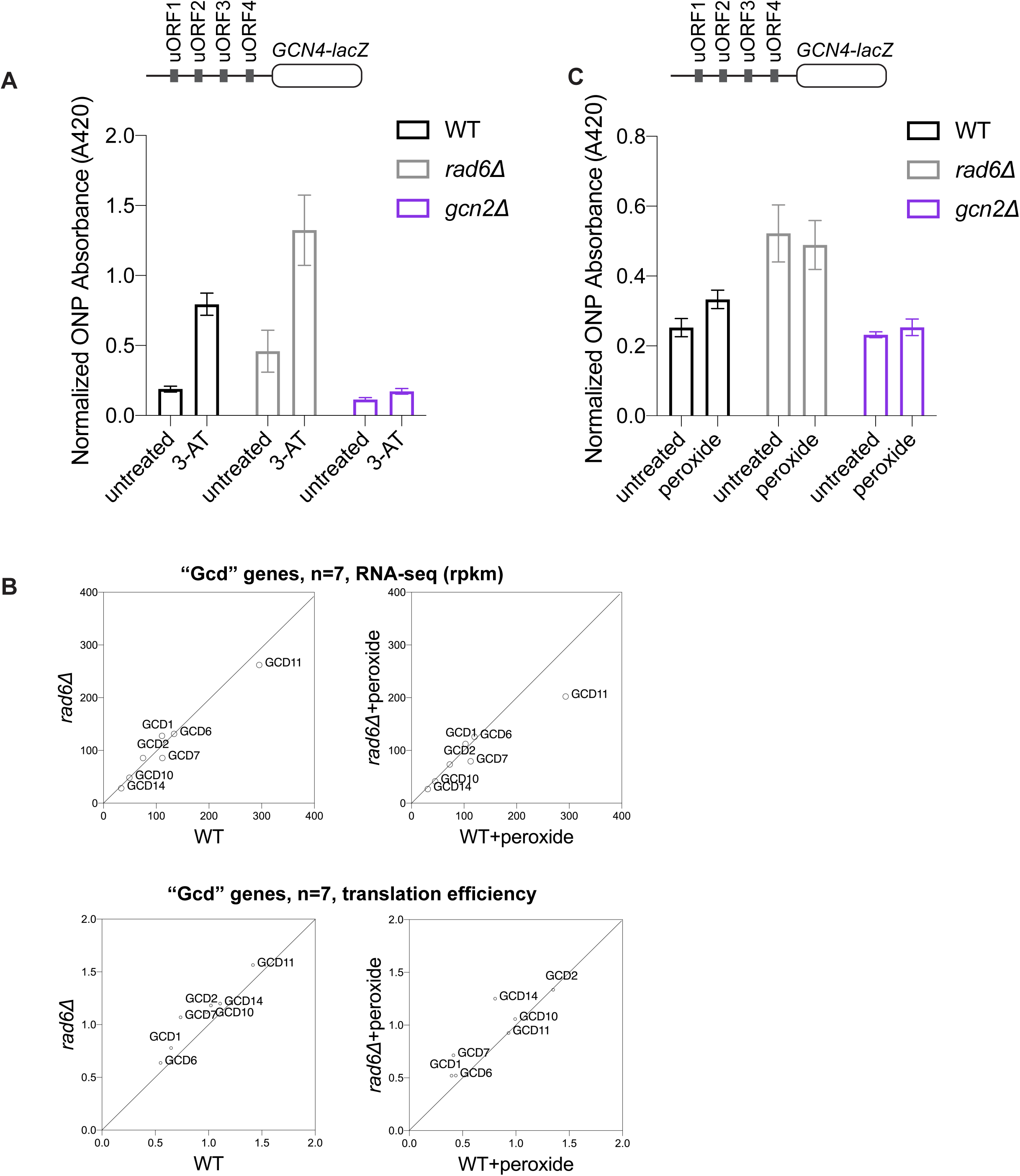
Supporting *GCN4-lacZ* reporter experiments and analysis of expression of *GCD* genes. Related to Figure 6. **(A)** *GCN4-lacZ* assay in the presence of 3-AT shows that Gcn4 translation is induced with 3-AT treatment, and this is dependent on the presence of Gcn2. These positive controls show the proper functioning of the reporter in these cells and also show that Gcn2 remains active in *rad6*Δ cells. **(B)** RNA-seq (top) and translation efficiency (bottom) data corresponding to mRNAs that encode “Gcd” genes indicate that the expression of these genes does not change in the cells lacking Rad6 ± peroxide. **(C)** *GCN4-lacZ* assay performed in the presence of peroxide. The data show that Gcn4 translation is higher in *rad6*Δ cells compared to WT, with and without peroxide.

**Table S1.** Yeast strains used in this study.

| Yeast name | Genotype | Source |
| --- | --- | --- |
| SUB280 WT | *MATa lys2-801 leu2-3,112 ura3-52 his3- Δ200 trp1-1[am] ubi1-Δ1::TRP1 ubi2-Δ2::ura3 ubi3-Δub-2 ubi4-Δ2::LEU2 [pUB39 Ub, LYS2][pUB100 Ubi1 tail, HIS3]* | (Finley et al., 1994) |
| SUB280 *rad6Δ* | *MATa lys2-801 leu2-3,112 ura3-52 his3- Δ200 trp1-1[am] ubi1-Δ1::TRP1 ubi2-Δ2::ura3 ubi3-Δub-2 ubi4-Δ2::LEU2 [pUB39 Ub, LYS2][pUB100 Ubi1 tail, HIS3] rad6:kanMX4* | (Silva et al., 2015) |
| SUB280 *hel2Δ* | *MATa lys2-801 leu2-3,112 ura3-52 his3- Δ200 trp1-1[am] ubi1-Δ1::TRP1 ubi2-Δ2::ura3 ubi3-Δub-2 ubi4-Δ2::LEU2 [pUB39 Ub, LYS2][pUB100 Ubi1 tail, HIS3] hel2:kanMX4* | (Back et al., 2019) |
| SUB280 *rad6Δ RAD6* | *MATa lys2-801 leu2-3,112 ura3-52 his3- Δ200 trp1-1[am] ubi1-Δ1::TRP1 ubi2-Δ2::ura3 ubi3-Δub-2 ubi4-Δ2::LEU2 [pUB39 Ub, LYS2][pUB100 Ubi1 tail, HIS3] [pYES RAD6-HA, URA3]* | (Simoes et al., 2022) |
| SUB280 *rad6Δ RAD6 (C88A)* | *MATa lys2-801 leu2-3,112 ura3-52 his3- Δ200 trp1-1[am] ubi1-Δ1::TRP1 ubi2-Δ2::ura3 ubi3-Δub-2 ubi4-Δ2::LEU2 [pUB39 Ub, LYS2][pUB100 Ubi1 tail, HIS3] [pYES RAD6(C88A)-HA, URA3]* | (Simoes et al., 2022) |
| S288C WT | *MATa SUC2 gal2 mal2 mel flo1 flo8-1 hap1 ho bio1 bio6* | (Mortimer and Johnston, 1986) |
| S288C *rad6Δ* | *MATa SUC2 gal2 mal2 mel flo1 flo8-1 hap1 ho bio1 bio6 rad6:kanMX4* | (Winzeler et al., 1999) |

**Table S2.**
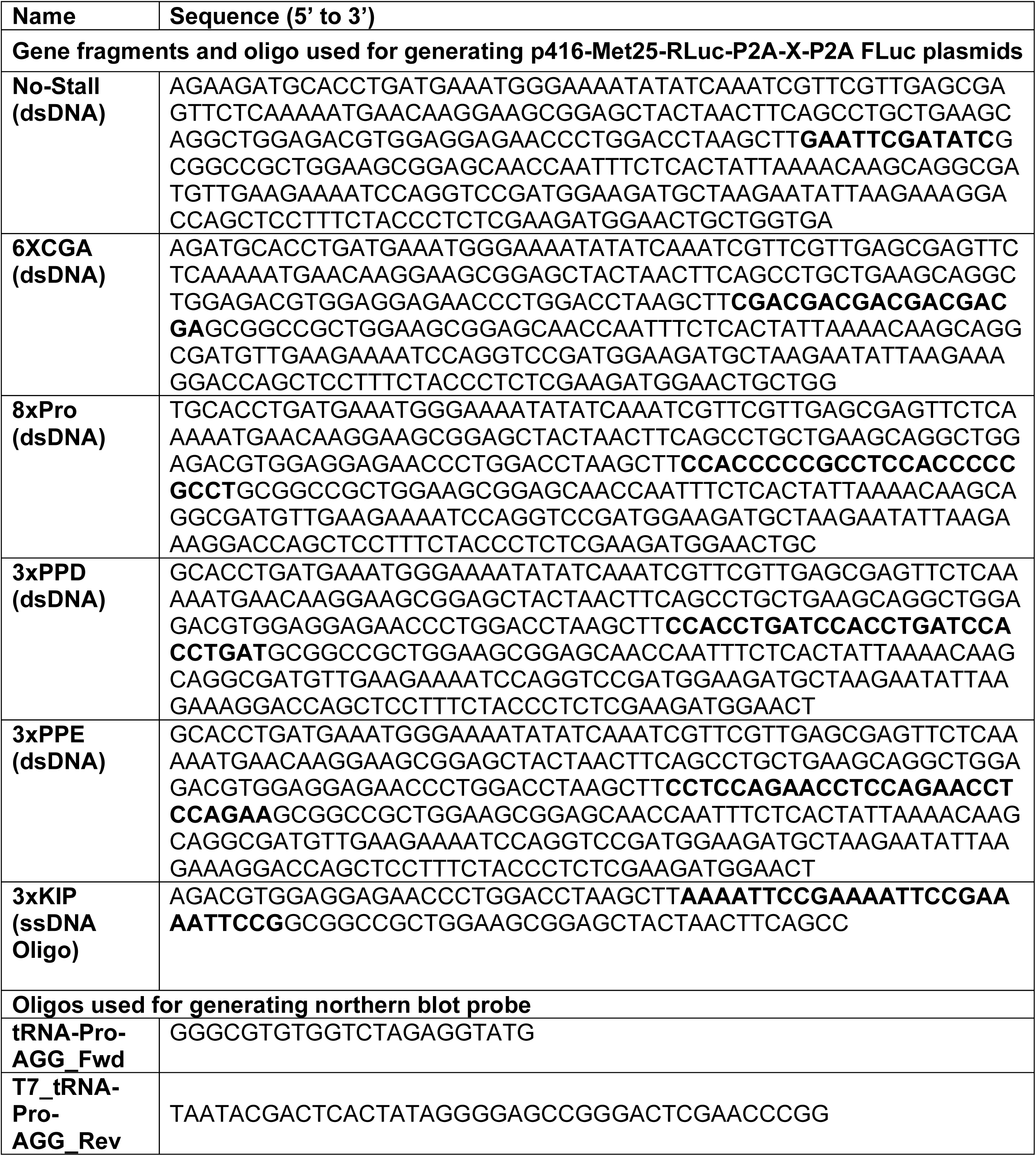
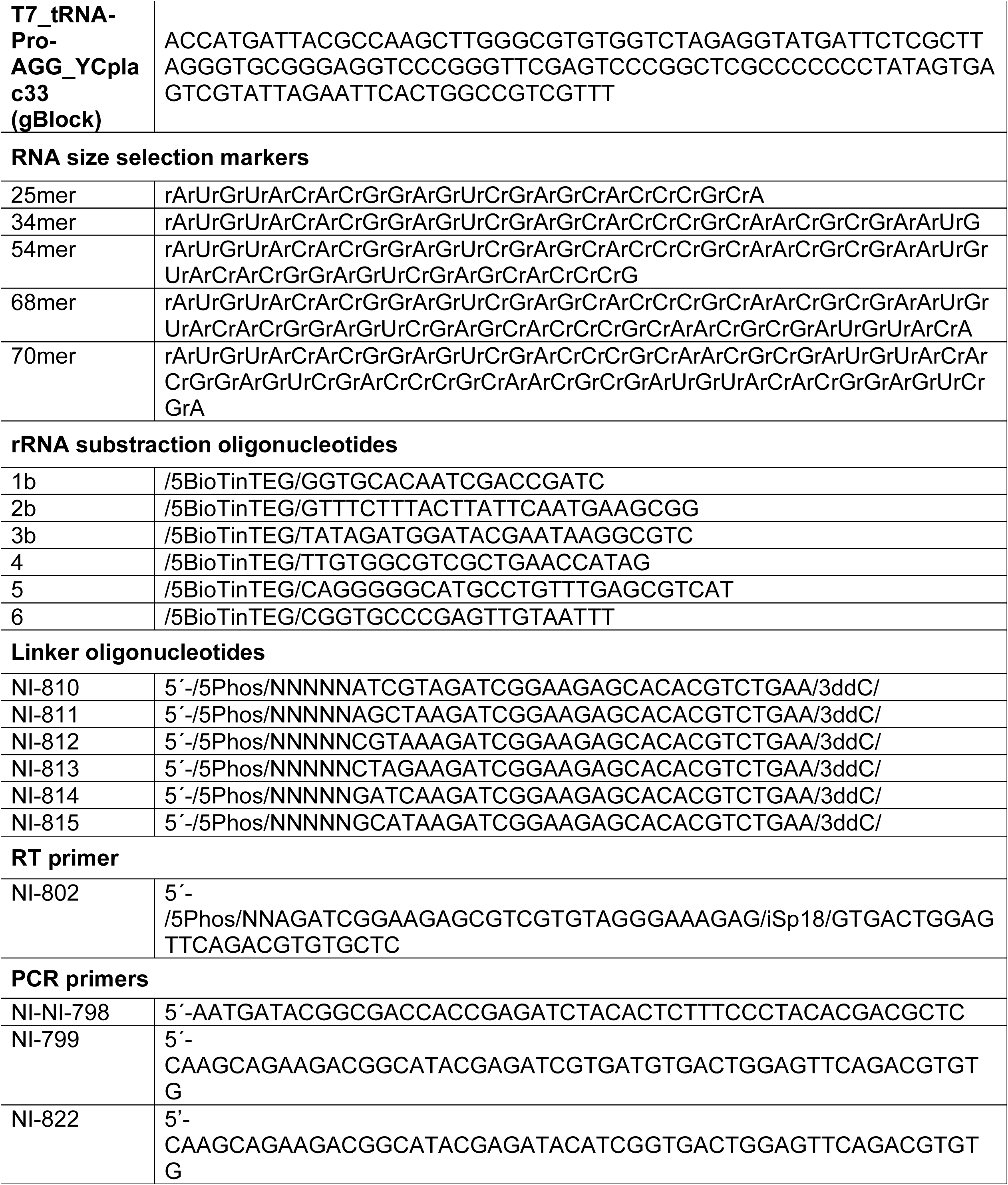
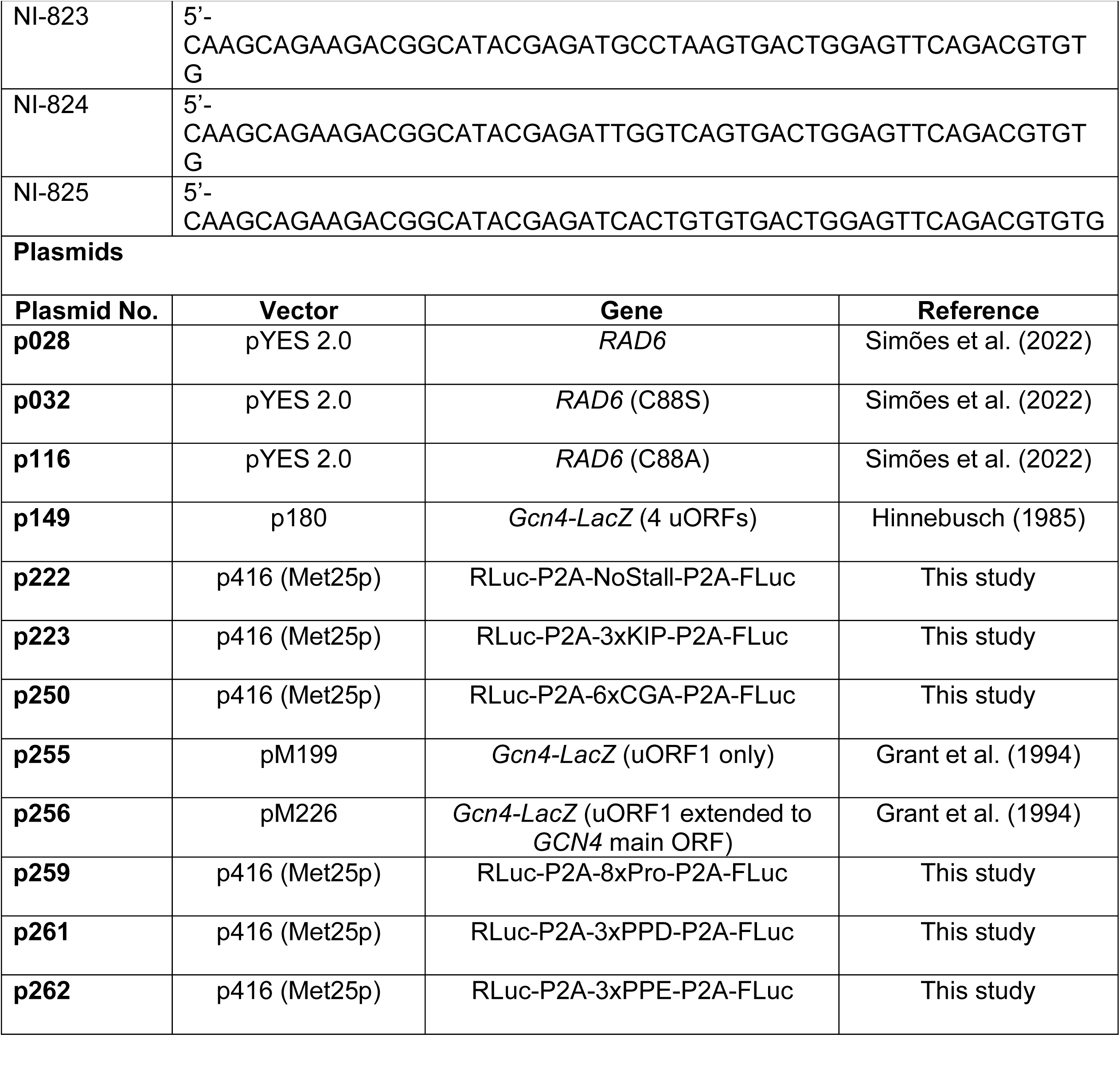
Oligonucleotides and plasmids used in this study.

**Table S3.** Ribosome profiling statistics.

| Sample name | Description | Data type | Reads with linker | Not aligned to non-coding RNA | Reads without PCR duplicates | Aligned to coding regions and splice junctions |
| --- | --- | --- | --- | --- | --- | --- |
| SM061F | SUB280 WT_rep1 | Ribo-seq | 88,465,576 | 32,310,998 | 21,072,905 | 16,214,578 |
| SM071F | SUB280 WT_rep2 | Ribo-seq | 81,035,725 | 41,623,662 | 29,668,687 | 24,022,020 |
| SM062F | SUB280 WT+peroxide_rep1 | Ribo-seq | 106,692,905 | 28,338,349 | 18,298,626 | 13,749,540 |
| SM072F | SUB280 WT+peroxide_rep2 | Ribo-seq | 78,751,014 | 26,997,840 | 19,255,686 | 15,370,434 |
| SM065F | SUB280 *rad6*Δ_rep1 | Ribo-seq | 91,450,355 | 32,530,594 | 13,938,148 | 10,430,067 |
| SM075F | SUB280 *rad6*Δ_rep2 | Ribo-seq | 102,956,589 | 43,203,482 | 30,983,491 | 24,965,084 |
| SM066F | SUB280 *rad6*Δ+peroxide_rep1 | Ribo-seq | 83,007,104 | 20,487,638 | 8,791,438 | 6,516,848 |
| SM076F | SUB280 *rad6*Δ+peroxide_rep2 | Ribo-seq | 83,266,747 | 28,388,419 | 20,644,308 | 17,042,575 |
| SM099F | SUB280 *hel2*Δ | Ribo-seq | 51,976,975 | 19,589,386 | 17,383,785 | 13,504,129 |
| SM100F | SUB280 *hel2*Δ+peroxide | Ribo-seq | 39,728,199 | 17,108,518 | 15,205,421 | 10,719,351 |
| SM103F | SUB280 *rad6*Δ *RAD6* | Ribo-seq | 40,183,394 | 26,152,266 | 20,545,187 | 15,387,454 |
| SM104F | SUB280 *rad6*Δ *RAD6*+peroxide | Ribo-seq | 38,330,665 | 16,932,848 | 13,310,052 | 10,124,867 |
| SM105F | SUB280 *rad6*Δ *RAD6* (C88A) | Ribo-seq | 73,490,623 | 39,788,606 | 30,908,395 | 24,485,828 |
| SM106F | SUB280 *rad6*Δ *RAD6* (C88A)+peroxide | Ribo-seq | 46,123,764 | 21,249,347 | 16,905,721 | 13,384,694 |
| SM107F | S288C WT | Ribo-seq | 43,906,581 | 18,322,535 | 15,662,623 | 11,286,980 |
| SM108F | S288C WT+peroxide | Ribo-seq | 46,024,866 | 10,218,846 | 8,698,065 | 6,022,004 |
| SM109F | S288C *rad6Δ* | Ribo-seq | 76,335,435 | 27,695,810 | 23,610,645 | 17,854,558 |
| SM110F | S288C *rad6Δ*+peroxide | Ribo-seq | 10,886,937 | 3,519,792 | 3,057,926 | 2,286,614 |
| SM061Fd | SUB280 WT_rep1 | Disome-seq | 7,761,659 | 5,475,884 | 3,524,369 | 964,664 |
| SM071Fd | SUB280 WT_rep2 | Disome-seq | 51,257,892 | 30,340,944 | 4,043,876 | 2,476,773 |
| SM062Fd | SUB280 WT+peroxide_rep1 | Disome-seq | 6,670,378 | 5,794,983 | 3,619,232 | 1,318,655 |
| SM072Fd | SUB280 WT+peroxide_rep2 | Disome-seq | 31,682,362 | 37,844,878 | 4,942,963 | 3,420,958 |
| SM065F | SUB280 *rad6Δ* | Disome-seq | 6,889,699 | 7,241,843 | 2,174,826 | 1,285,476 |
| SM066F | SUB280 *rad6Δ*+peroxide | Disome-seq | 4,596,077 | 7,346,885 | 2,800,625 | 733,394 |
| SM061M | SUB280 WT_rep1 | RNA-seq | 72,941,175 | 34,744,878 | 24,913,881 | 16,298,506 |
| SM071M | SUB280 WT_rep2 | RNA-seq | 68,143,036 | 26,850,953 | 20,027,139 | 11,889,363 |
| SM062M | SUB280 WT+peroxide_rep1 | RNA-seq | 61,818,591 | 26,895,510 | 19,329,586 | 12,833,259 |
| SM072M | SUB280 WT+peroxide_rep2 | RNA-seq | 73,905,111 | 26,350,153 | 19,532,817 | 11,208,925 |
| SM065M | SUB280 *rad6Δ*_rep1 | RNA-seq | 100,532,895 | 34,833,857 | 24,941,388 | 15,815,962 |
| SM075M | SUB280 *rad6Δ*_rep2 | RNA-seq | 68,204,722 | 23,096,946 | 16,515,446 | 8,793,155 |
| SM066M | SUB280 *rad6Δ*+peroxide_rep1 | RNA-seq | 146,467,948 | 40,302,699 | 29,535,984 | 15,823,150 |
| SM076M | SUB280 *rad6Δ*+peroxide_rep2 | RNA-seq | 81,834,980 | 20,039,453 | 14,121,673 | 7,496,153 |

## Notes

### Competing Interest Statement

The authors have declared no competing interest.

## Bibliography

Advani, V.M., and Ivanov, P. (2019). Translational Control under Stress: Reshaping the Translatome. Bioessays 41, e1900009.

Arima, Y., Nishigori, C., Takeuchi, T., Oka, S., Morimoto, K., Utani, A., and Miyachi, Y. (2006). 4-Nitroquinoline 1-oxide forms 8-hydroxydeoxyguanosine in human fibroblasts through reactive oxygen species. Toxicol Sci 91, 382–392.

Back, S., Gorman, A.W., Vogel, C., and Silva, G.M. (2019). Site-Specific K63 Ubiquitinomics Provides Insights into Translation Regulation under Stress. J Proteome Res 18, 309–318.

Beckhouse, A.G., Grant, C.M., Rogers, P.J., Dawes, I.W., and Higgins, V.J. (2008). The adaptive response of anaerobically grown Saccharomyces cerevisiae to hydrogen peroxide is mediated by the Yap1 and Skn7 transcription factors. FEMS Yeast Res 8, 1214–1222.

Ben Rejeb, K., Abdelly, C., and Savoure, A. (2014). How reactive oxygen species and proline face stress together. Plant Physiol Biochem 80, 278–284.

Brandman, O., Stewart-Ornstein, J., Wong, D., Larson, A., Williams, C.C., Li, G.W., Zhou, S., King, D., Shen, P.S., Weibezahn, J., et al. (2012). A ribosome-bound quality control complex triggers degradation of nascent peptides and signals translation stress. Cell 151, 1042–1054.

Bresson, S., Shchepachev, V., Spanos, C., Turowski, T.W., Rappsilber, J., and Tollervey, D. (2020). Stress-Induced Translation Inhibition through Rapid Displacement of Scanning Initiation Factors. Mol Cell 80, 470–484 e478.

Bruinsma, C.F., Savelberg, S.M., Kool, M.J., Jolfaei, M.A., Van Woerden, G.M., Baarends, W.M., and Elgersma, Y. (2016). An essential role for UBE2A/HR6A in learning and memory and mGLUR-dependent long-term depression. Hum Mol Genet 25, 1–8.

Chan, C.T., Pang, Y.L., Deng, W., Babu, I.R., Dyavaiah, M., Begley, T.J., and Dedon, P.C. (2012). Reprogramming of tRNA modifications controls the oxidative stress response by codon-biased translation of proteins. Nat Commun 3, 937.

Cherkasova, V.A., and Hinnebusch, A.G. (2003). Translational control by TOR and TAP42 through dephosphorylation of eIF2αlpha kinase GCN2. Genes Dev 17, 859–872.

Czeschik, J.C., Bauer, P., Buiting, K., Dufke, C., Guillen-Navarro, E., Johnson, D.S., Koehler, U., Lopez-Gonzalez, V., Ludecke, H.J., Male, A., et al. (2013). X-linked intellectual disability type Nascimento is a clinically distinct, probably underdiagnosed entity. Orphanet J Rare Dis 8, 146.

Dougherty, S.E., Maduka, A.O., Inada, T., and Silva, G.M. (2020). Expanding Role of Ubiquitin in Translational Control. Int J Mol Sci 21.

Finkel, T., and Holbrook, N.J. (2000). Oxidants, oxidative stress and the biology of ageing. Nature 408, 239–247.

Finley, D., Sadis, S., Monia, B.P., Boucher, P., Ecker, D.J., Crooke, S.T., and Chau, V. (1994). Inhibition of proteolysis and cell cycle progression in a multiubiquitination-deficient yeast mutant. Mol Cell Biol 14, 5501–5509.

Gasch, A.P., Spellman, P.T., Kao, C.M., Carmel-Harel, O., Eisen, M.B., Storz, G., Botstein, D., and Brown, P.O. (2000). Genomic expression programs in the response of yeast cells to environmental changes. Mol Biol Cell 11, 4241–4257.

Gerashchenko, M.V., and Gladyshev, V.N. (2014). Translation inhibitors cause abnormalities in ribosome profiling experiments. Nucleic Acids Res 42, e134.

Gerashchenko, M.V., Lobanov, A.V., and Gladyshev, V.N. (2012). Genome-wide ribosome profiling reveals complex translational regulation in response to oxidative stress. Proc Natl Acad Sci U S A 109, 17394–17399.

Grant, C.M. (2011). Regulation of translation by hydrogen peroxide. Antioxid Redox Signal 15, 191–203.

Gu, C., Begley, T.J., and Dedon, P.C. (2014). tRNA modifications regulate translation during cellular stress. FEBS Lett 588, 4287–4296.

Guydosh, N.R., and Green, R. (2014). Dom34 rescues ribosomes in 3’ untranslated regions. Cell 156, 950–962.

Haddad, D.M., Vilain, S., Vos, M., Esposito, G., Matta, S., Kalscheuer, V.M., Craessaerts, K., Leyssen, M., Nascimento, R.M., Vianna-Morgante, A.M., et al. (2013). Mutations in the intellectual disability gene Ube2a cause neuronal dysfunction and impair parkin-dependent mitophagy. Mol Cell 50, 831–843.

Harding, H.P., Novoa, I., Zhang, Y., Zeng, H., Wek, R., Schapira, M., and Ron, D. (2000). Regulated translation initiation controls stress-induced gene expression in mammalian cells. Mol Cell 6, 1099–1108.

Hinnebusch, A.G. (2005). Translational regulation of GCN4 and the general amino acid control of yeast. Annu Rev Microbiol 59, 407–450.

Houston, L., Platten, E.M., Connelly, S.M., Wang, J., and Grayhack, E.J. (2022). Frameshifting at collided ribosomes is modulated by elongation factor eEF3 and by integrated stress response regulators Gcn1 and Gcn20. RNA 28, 320–339.

Ikeuchi, K., Tesina, P., Matsuo, Y., Sugiyama, T., Cheng, J., Saeki, Y., Tanaka, K., Becker, T., Beckmann, R., and Inada, T. (2019). Collided ribosomes form a unique structural interface to induce Hel2-driven quality control pathways. EMBO J 38.

Ingolia, N.T., Ghaemmaghami, S., Newman, J.R., and Weissman, J.S. (2009). Genome-wide analysis in vivo of translation with nucleotide resolution using ribosome profiling. Science 324, 218–223.

Ingolia, N.T., Hussmann, J.A., and Weissman, J.S. (2019). Ribosome Profiling: Global Views of Translation. Cold Spring Harb Perspect Biol 11.

Klopotowski, T., and Wiater, A. (1965). Synergism of aminotriazole and phosphate on the inhibition of yeast imidazole glycerol phosphate dehydratase. Arch Biochem Biophys 112, 562–566.

Kubota, H., Obata, T., Ota, K., Sasaki, T., and Ito, T. (2003). Rapamycin-induced translational derepression of GCN4 mRNA involves a novel mechanism for activation of the eIF2 alpha kinase GCN2. J Biol Chem 278, 20457–20460.

Langmead, B., Trapnell, C., Pop, M., and Salzberg, S.L. (2009). Ultrafast and memory-efficient alignment of short DNA sequences to the human genome. Genome Biol 10, R25.

Letzring, D.P., Wolf, A.S., Brule, C.E., and Grayhack, E.J. (2013). Translation of CGA codon repeats in yeast involves quality control components and ribosomal protein L1. RNA 19, 1208–1217.

Levings, D.C., Lacher, S.E., Palacios-Moreno, J., and Slattery, M. (2021). Transcriptional reprogramming by oxidative stress occurs within a predefined chromatin accessibility landscape. Free Radic Biol Med 171, 319–331.

Liang, X., Zhang, L., Natarajan, S.K., and Becker, D.F. (2013). Proline mechanisms of stress survival. Antioxid Redox Signal 19, 998–1011.

Liguori, I., Russo, G., Curcio, F., Bulli, G., Aran, L., Della-Morte, D., Gargiulo, G., Testa, G., Cacciatore, F., Bonaduce, D., et al. (2018). Oxidative stress, aging, and diseases. Clin Interv Aging 13, 757–772.

Love, M.I., Huber, W., and Anders, S. (2014). Moderated estimation of fold change and dispersion for RNA-seq data with DESeq2. Genome Biol 15, 550.

Ma, Q. (2010). Transcriptional responses to oxidative stress: pathological and toxicological implications. Pharmacol Ther 125, 376–393.

Marguerat, S., Lawler, K., Brazma, A., and Bahler, J. (2014). Contributions of transcription and mRNA decay to gene expression dynamics of fission yeast in response to oxidative stress. RNA Biol 11, 702–714.

Martin, M. (2011). Cutadapt removes adapter sequences from high-throughput sequencing reads. 2011 *17*, 3.

Mascarenhas, C., Edwards-Ingram, L.C., Zeef, L., Shenton, D., Ashe, M.P., and Grant, C.M. (2008). Gcn4 is required for the response to peroxide stress in the yeast Saccharomyces cerevisiae. Mol Biol Cell 19, 2995–3007.

Matsuo, Y., Ikeuchi, K., Saeki, Y., Iwasaki, S., Schmidt, C., Udagawa, T., Sato, F., Tsuchiya, H., Becker, T., Tanaka, K., et al. (2017). Ubiquitination of stalled ribosome triggers ribosome-associated quality control. Nat Commun 8, 159.

Matsuo, Y., Tesina, P., Nakajima, S., Mizuno, M., Endo, A., Buschauer, R., Cheng, J., Shounai, O., Ikeuchi, K., Saeki, Y., et al. (2020). RQT complex dissociates ribosomes collided on endogenous RQC substrate SDD1. Nat Struct Mol Biol 27, 323–332.

McGlincy, N.J., and Ingolia, N.T. (2017). Transcriptome-wide measurement of translation by ribosome profiling. Methods 126, 112–129.

Meydan, S., and Guydosh, N.R. (2020). Disome and Trisome Profiling Reveal Genome-wide Targets of Ribosome Quality Control. Mol Cell 79, 588–602 e586.

Mi, H., Muruganujan, A., and Thomas, P.D. (2013). PANTHER in 2013: modeling the evolution of gene function, and other gene attributes, in the context of phylogenetic trees. Nucleic Acids Res 41, D377–386.

Mortimer, R.K., and Johnston, J.R. (1986). Genealogy of principal strains of the yeast genetic stock center. Genetics 113, 35–43.

Nascimento, R.M., Otto, P.A., de Brouwer, A.P., and Vianna-Morgante, A.M. (2006). UBE2A, which encodes a ubiquitin-conjugating enzyme, is mutated in a novel X-linked mental retardation syndrome. Am J Hum Genet 79, 549–555.

Natarajan, K., Meyer, M.R., Jackson, B.M., Slade, D., Roberts, C., Hinnebusch, A.G., and Marton, M.J. (2001). Transcriptional profiling shows that Gcn4p is a master regulator of gene expression during amino acid starvation in yeast. Mol Cell Biol 21, 4347–4368.

Ng, P.C., Wong, E.D., MacPherson, K.A., Aleksander, S., Argasinska, J., Dunn, B., Nash, R.S., Skrzypek, M.S., Gondwe, F., Jha, S., et al. (2020). Transcriptome visualization and data availability at the Saccharomyces Genome Database. Nucleic Acids Res 48, D743–D748.

O’Shea, J.P., Chou, M.F., Quader, S.A., Ryan, J.K., Church, G.M., and Schwartz, D. (2013). pLogo: a probabilistic approach to visualizing sequence motifs. Nat Methods 10, 1211–1212.

Pelechano, V., Wei, W., Jakob, P., and Steinmetz, L.M. (2014). Genome-wide identification of transcript start and end sites by transcript isoform sequencing. Nat Protoc 9, 1740–1759.

Picazo, C., and Molin, M. (2021). Impact of Hydrogen Peroxide on Protein Synthesis in Yeast. Antioxidants (Basel) 10.

Qiu, H., Chereji, R.V., Hu, C., Cole, H.A., Rawal, Y., Clark, D.J., and Hinnebusch, A.G. (2016). Genome-wide cooperation by HAT Gcn5, remodeler SWI/SNF, and chaperone Ydj1 in promoter nucleosome eviction and transcriptional activation. Genome Res 26, 211–225.

Rawal, Y., Chereji, R.V., Valabhoju, V., Qiu, H., Ocampo, J., Clark, D.J., and Hinnebusch, A.G. (2018). Gcn4 Binding in Coding Regions Can Activate Internal and Canonical 5’ Promoters in Yeast. Mol Cell 70, 297–311 e294.

Rubio, A., Ghosh, S., Mulleder, M., Ralser, M., and Mata, J. (2021). Ribosome profiling reveals ribosome stalling on tryptophan codons and ribosome queuing upon oxidative stress in fission yeast. Nucleic Acids Res 49, 383–399.

Saint, M., Sawhney, S., Sinha, I., Singh, R.P., Dahiya, R., Thakur, A., Siddharthan, R., and Natarajan, K. (2014). The TAF9 C-terminal conserved region domain is required for SAGA and TFIID promoter occupancy to promote transcriptional activation. Mol Cell Biol 34, 1547–1563.

Sharifi-Rad, M., Anil Kumar, N.V., Zucca, P., Varoni, E.M., Dini, L., Panzarini, E., Rajkovic, J., Tsouh Fokou, P.V., Azzini, E., Peluso, I., et al. (2020). Lifestyle, Oxidative Stress, and Antioxidants: Back and Forth in the Pathophysiology of Chronic Diseases. Front Physiol 11, 694.

Shenton, D., Smirnova, J.B., Selley, J.N., Carroll, K., Hubbard, S.J., Pavitt, G.D., Ashe, M.P., and Grant, C.M. (2006). Global translational responses to oxidative stress impact upon multiple levels of protein synthesis. J Biol Chem 281, 29011–29021.

Silva, G.M., Finley, D., and Vogel, C. (2015). K63 polyubiquitination is a new modulator of the oxidative stress response. Nat Struct Mol Biol 22, 116–123.

Simoes, V., Cizubu, B.K., Harley, L., Zhou, Y., Pajak, J., Snyder, N.A., Bouvette, J., Borgnia, M.J., Arya, G., Bartesaghi, A., et al. (2022). Redox-sensitive E2 Rad6 controls cellular response to oxidative stress via K63-linked ubiquitination of ribosomes. Cell Rep 39, 110860.

Szabados, L., and Savoure, A. (2010). Proline: a multifunctional amino acid. Trends Plant Sci 15, 89–97.

Takagi, H. (2008). Proline as a stress protectant in yeast: physiological functions, metabolic regulations, and biotechnological applications. Appl Microbiol Biotechnol 81, 211–223.

Thomas, P.D., Ebert, D., Muruganujan, A., Mushayahama, T., Albou, L.P., and Mi, H. (2022). PANTHER: Making genome-scale phylogenetics accessible to all. Protein Sci 31, 8–22.

Thompson, D.M., Lu, C., Green, P.J., and Parker, R. (2008). tRNA cleavage is a conserved response to oxidative stress in eukaryotes. RNA 14, 2095–2103.

Wickham, H. (2016). ggplot2: Elegant Graphics for Data Analysis (Springer-Verlag New York).

Winzeler, E.A., Shoemaker, D.D., Astromoff, A., Liang, H., Anderson, K., Andre, B., Bangham, R., Benito, R., Boeke, J.D., Bussey, H., et al. (1999). Functional characterization of the S. cerevisiae genome by gene deletion and parallel analysis. Science 285, 901–906.

Wu, C.C., Zinshteyn, B., Wehner, K.A., and Green, R. (2019). High-Resolution Ribosome Profiling Defines Discrete Ribosome Elongation States and Translational Regulation during Cellular Stress. Mol Cell 73, 959–970 e955.

Yan, L.L., Simms, C.L., McLoughlin, F., Vierstra, R.D., and Zaher, H.S. (2019). Oxidation and alkylation stresses activate ribosome-quality control. Nat Commun 10, 5611.

Yan, L.L., and Zaher, H.S. (2021). Ribosome quality control antagonizes the activation of the integrated stress response on colliding ribosomes. Mol Cell 81, 614–628 e614.

Young, M.J., and Court, D.A. (2008). Effects of the S288c genetic background and common auxotrophic markers on mitochondrial DNA function in Saccharomyces cerevisiae. Yeast 25, 903–912.

Zhao, T., Chen, Y.M., Li, Y., Wang, J., Chen, S., Gao, N., and Qian, W. (2021). Disome-seq reveals widespread ribosome collisions that promote cotranslational protein folding. Genome Biol 22, 16.

Zhou, Y., Kastritis, P.L., Dougherty, S.E., Bouvette, J., Hsu, A.L., Burbaum, L., Mosalaganti, S., Pfeffer, S., Hagen, W.J.H., Forster, F., et al. (2020). Structural impact of K63 ubiquitin on yeast translocating ribosomes under oxidative stress. Proc Natl Acad Sci U S A 117, 22157–22166.

